# Metabolic and imaging phenotypes associated with *RB1* and *TP53* loss in prostate cancer

**DOI:** 10.1101/2023.11.15.567250

**Authors:** Fahim Ahmad, Margaret White, Kazutoshi Yamamoto, Daniel R. Crooks, Supreet Agarwal, Ye Yang, Brian Capaldo, Sonam Raj, Aian Neil Alilin, Anita Ton, Stephen Adler, Jurgen Seidel, Colleen Olkowski, Murali Krishna Cherukuri, Peter L Choyke, Kathleen Kelly, Jeffrey R. Brender

## Abstract

Advanced prostate cancer is treated with androgen receptor (AR) signaling inhibitors, which are initially effective, but most patients eventually develop resistance and progress to castrate-resistant prostate cancer (CRPC). Loss of RB1 in CRPC tumors is correlated with rapid progression and poor patient survival and, in combination with TP53 loss, predisposes patients to the development of transitional neuroendocrine prostate cancer (NEPC). Although progressive CRPC is clinically associated with higher 18FDG-PET SUVmax values, it is unknown whether inactivation of RB1 and/or TP53 is a driver of increased glucose import. Using a cohort of patient-derived xenograft (PDX)-derived CRPC organoids, we found that NEPC could not be conclusively distinguished from adenocarcinoma by 18FDG uptake alone, and PSMA protein levels did not correlate with cancer phenotype or 18FDG uptake. Castration-resistant models showed higher 18FDG uptake, but lower pyruvate-to-lactate conversion compared to their castration-sensitive counterparts. In parallel studies using castration-sensitive prostate cancer models, RB1/TP53 knockdown did not affect 18FDG uptake, but increased basal respiration and glycolytic activity, with combined depletion leading to glucose diversion into glycogenesis. These metabolic changes were reflected in increased lactate dehydrogenase flux detected by 13C-hyperpolarized magnetic resonance spectroscopy upon RB1 loss, but not in 18FDG uptake. The metabolic heterogeneity revealed here suggests that a multimodal molecular imaging approach can improve tumor characterization, potentially leading to a better prognosis in cancer treatment.

## 1. Introduction

Oncogenic transformation and progression is often associated with metabolic reprogramming and this association can be exploited for molecular imaging, most notably by FDG-PET. Prostate cancer is of great interest from a metabolic perspective due to the unusual metabolism of the prostate and the extent of metabolic reprogramming during oncogenic transformation. In contrast to most cells that oxidize citrate through the Krebs cycle, benign prostate epithelial cells are net citrate producers. [1,2] While the role of citrate in seminal fluid is not fully understood, it is of significant enough importance in fertilization that drastic and metabolically expensive adaptions have evolved to secrete it at extraordinarily high levels,[3] which is achieved by diverting citrate from the TCA cycle. As a result, prostate epithelial cells rely on alternative metabolic pathways for survival, primarily high levels of aerobic glycolysis, sacrificing metabolic efficiency for citrate production.(3)

This metabolic shift is primarily under the control of the Androgen Receptor (AR), [4,5] which regulates multiple aspects of metabolism through the AMPK-PFK pathway to increase glucose uptake, glycolysis, lipogenesis, and fatty acid oxidation. [4,6,7] Since prostate cancer typically develops in the same epithelial cells responsible for citrate production, the energetically expensive diversion of the TCA cycle poses an obstacle to proliferation. During oncogenic transformation, prostate cancer cells reverse this phenotype and adopt a zinc-wasting, citrate-oxidizing phenotype,(3) building on the same underlying circuitry as the basal metabolism but directing it towards a different purpose.

Aggressive and/or recurrent prostate cancer is treated by inhibiting androgen receptor (AR) signaling, a pathway required by most tumor cells for growth and viability.[8] Although treatment is usually initially effective, resistance to AR inhibition almost always develops, leading to the emergence of castration-resistant prostate cancer (CRPC). While most CRPC circumvents AR pathway inhibition by upregulating AR signaling either directly or indirectly, approximately 15-20% of CRPC patients lose dependence upon AR signaling.[8,9] The de-differentiated neuroendocrine prostate cancer (NEPC) tumors that emerge from CRPC in particular are notable for diminished AR signaling and an unfavorable prognosis.[10,11] The loss of AR signaling in NEPC is accompanied by profound metabolic reprogramming, including increased glycolysis and altered glucose utilization, which are hallmarks of aggressive disease progression. However, the complexity of AR signal dependent metabolic changes has made it difficult to obtain a complete molecular characterization of prostate cancer metabolism. AR signaling is dependent on a number of downstream effectors which, depending on the environmental context, can either amplify or reverse the signal. While metabolic differences in individual tumors likely contribute to the efficacy of some cancer treatments, CRPC is highly heterogeneous,[12,13] and the range of metabolic reprogramming as well as association with specific phenotypic and genotypic variations are unknown.

Analysis of recurrent CRPC genomic alterations relative to clinical outcome has identified *RB1* homozygous deletion, occurring in about 10% of CRPC cases, as singularly associated with poor survival.[14] Homozygous alterations in *RB1* and *TP53* are near universal in NEPC, [15,16] along with a loss of AR expression which leads to increased cellular plasticity and differentiation along a neuroendocrine lineage.[17] However, *RB1/TP53* loss is not unique to NEPC and there are androgen receptor positive adenocarcinomas (ARPCs) that harbor alterations in *RB1* and *TP53*.[18] ARPCs with alterations in *RB1* continue to signal through the AR but show changes in proliferation and DNA repair pathways [18] While the loss of *RB1* alone is not determinative, it greatly increases the likelihood of a transition to NEPC, making the detection of *RB1* loss in CRPC a critical factor with significant implications for tailoring treatment strategies and patient monitoring.[14]

The effect of *RB1* loss is known to be strongly species and context dependent, having almost no pathological effect in some tissues but driving carcinogenesis in others.[19] In prostate cancer, the effect of *RB1* loss seems to be mediated by *TP53*, with dual *RB1/TP53* loss associated with a decrease in AR signaling not seen with the inactivation of either gene alone, activating a transcriptional program that creates a stem cell-like phenotype that is resistant to diverse forms of chemotherapy.[18] Like RB1, *TP53* is heavily involved in the regulation of metabolism, suppressing glucose metabolism by downregulation of hexokinase 2 and reducing transcription of the glucose transporter GLUT1 and preventing translocation to the plasma membrane.[20] Inactivation of *TP53* therefore contributes to a glycolytic phenotype similar to *RB1* but through an alternate signaling pathway. Because signaling from *RB1* is separate and distinct from *TP5*3 and may act at cross-purposes,[18,21] it is unknown whether the increase in glycolysis seen in some CRPC tumors and nearly all NEPC tumors is a direct result of in the dual inactivation of *RB1* and *TP53*.

Clinically, neuroendocrine prostate cancer (NEPC) and aggressive CRPC metastases are characterized by increased FDG uptake on ^18^FDG-PET, which provides a convenient method for monitoring disease progression and treatment response. However, the molecular basis for this increased FDG uptake remains poorly understood. Given that both *RB1/TP53* loss and increased FDG uptake are characteristic features of NEPC, a critical question emerges: Is *RB1/TP53* loss sufficient to drive the metabolic reprogramming—particularly increased glucose uptake—observed in NEPC, even in the context of maintained AR signaling and castration sensitivity? Understanding this relationship would provide insight into the mechanisms underlying the metabolic transformation during prostate cancer progression and potentially identify early markers of aggressive disease.

Using an extensive cohort of newly developed clinically heterogeneous organoids[22] to investigate ^18^FDG uptake and conversion of pyruvate to lactate on hyperpolarized MRI (HP-MRI) relative to established CRPC phenotypes and genotype., we show here that NEPC cannot be conclusively distinguished from adenocarcinoma by ^18^FDG uptake alone, and PSMA protein levels do not correlate with cancer phenotype or ^18^FDG uptake in our models. In the limited cell lines where a direct comparison was possible, castration-resistant PDX models exhibited higher 18FDG uptake compared to their castration-sensitive counterparts but showed lower pyruvate-to-lactate conversion. Because RB1/TP53 deficiency predicts poor survival and increased transformation to a NEPC phenotype, we further investigated the metabolic changes in genomically stable castration sensitive models following *RB1/TP53* loss. In these models, *RB1/TP53* knockdown did not affect 18FDG uptake in vivo or in vitro. However, RB1 loss significantly increased basal respiration, glycolytic activity, and lactate dehydrogenase (LDH) flux, with combined *RB1/TP53* depletion leading to diversion of glucose into glycogenesis and enhanced TCA cycle activity. These metabolic alterations were detectable by ^13^C-hyperpolarized MRI spectroscopy but not by ^18^FDG uptake. Overall, our data does not support the hypothesis that NEPC tumor cells are responsible for increased FDG uptake observed clinically, but alternatively, such uptake likely occurs in non-malignant cells in the microenvironment which are metabolically activated by NEPC tumor biology.

## 2. Materials and Methods

### 2.1 3D Organoid culture

LuCaP PDX tumors were maintained at the NCI under the NCI Animal Care and Use Committee approved protocol (LGCP-003-2-C) and validated using STR analysis by Laragen, Inc. For processing, tissue was cut into small pieces, 2 to 4 mm, with a scalpel blade. The tissue was then collected in advanced DMEM/F12 with 10 mmol/L HEPES and 2 mmol/L Glutamax, transferred to gentle MACS C tubes (Miltenyi Biotec: #130-096-334) and digested using the Tumor Dissociation Kit for human tissue (Miltenyi Biotec: #130-095-929) on a gentle MACS Octo Dissociator with heaters (Miltenyi Biotec: #130-096-427) using program 37C_h_TDK_2. Processed samples were centrifuged, resuspended, and passed through a 100-μm cell strainer as needed to eliminate macroscopic tissue pieces. For PDXs, cell pellets were resuspended in two to three volumes of ACK lysis buffer (Lonza: #10-548E) according to the manufacturers’ specifications, and washed cells were resuspended in media.

PDX tumors maintained at the NCI were passaged in NOD SCID gamma (NSG) mice. After processing as described above, cell suspensions (2 × 10^6^ cells) were mixed with growth factor-reduced phenol red-free Matrigel (BD Biosciences, #356231) and injected subcutaneously.

To culture PDX-derived cells as organoids, cell suspensions, processed as described above, were plated in 3D on a 10 cm dish and 6 and 96 well plates in 5% Matrigel. The growth medium formulation for PrENR was described by Drost and colleagues.[23] For passaging, organoids were incubated with dispase, dissolved in advanced DMEM/F12 to a final concentration of 1 mg/mL, and incubated for 2 h at 37℃. pelleted organoids were resuspended in prewarmed TrypLE Express (Thermo Fisher: #12605028) for approximately 5 min at 37℃ with periodic pipetting.

The resulting cell suspensions were diluted in advanced DMEM/F12, centrifuged, resuspended, and plated, as described previously.

### 2.2 2D Cell Culture

LNCaP cells were purchased from ATCC and cultured in RPMI with 10% FBS, as previously described.[24]

### 2.3 Lentiviral transduction

A total of 2 × 10^6^ cells were transduced with various lentiviruses, such as SMARTvector Inducible Human Non-targeting Control, *RB1*, *TP53* shRNA (Dharmacon Reagents), and pLX313-Renilla Luciferase (Addgene; #118016) by spin infection at 1200 rpm for 90 min at room temperature. The cells were incubated overnight in the presence of the virus. The cells were then collected and replated in 3D Matrigel, as described previously.[22] Further, cells were selected against the prescribed antibiotic with 2.5 μg/mL of puromycin and hygromycin, respectively, for stable cell lines.

### 2.4 *RB1* and/or *TP53* LNCaP CRISPR-Cas9 knock-out

For CRISPR-mediated *RB1* and *TP53* gene deletion, we used *RB* and *TP53* guide sequences AAGTGAACGACATCTCATCTAGG and TATCTGAGCAGCGCTCATGGTGG (g)RNA, respectively, designed by the Broad Institute sgRNA designer. Blasticidin-resistant Cas9 along with sgRNA and GFP plasmids were electroporated in LNCaP cells, and the cells were cultured for 72 h. After 72 h, single GFP-positive cells were sorted and grown as monoclones for *RB1, TP53* and *RB1/TP53* Crispr KOs. Since WB showed that the expression of *RB1* and *TP53* proteins was completely abolished in all clones, we used only clone C1 of RB1 and/or TP53 Crispr KOs in subsequent experiments.

### 2.5 Generation of *RB1*, *TP53* altered PDX system

*RB1*, *TP53* and *RB1/TP53* altered LuCap 167 PDX tumors were generated in 6-week-old male NODscid gamma (NSG) mice by subcutaneously injecting *RB1*, *TP53*, *RB1/TP53* altered organoids (2 × 10^6^ cells) mixed with growth factor– reduced, phenol red–free Matrigel (BD Biosciences: #356231) under an NCI Animal Care and Use Committee approved protocol (LGCP-003-2-C). As the tumor started to grow, the mice were randomized and placed in two different groups [*RB1*, *TP53*, or *RB1/TP53* (control) and (depleted)]. Mice were given an IP injection of 100 ml of saline (control) or 100 ml of 0.2 mg/ml doxycycline prepared in saline, twice weekly until the tumor reaches 1-1.2 cm in diameter. The tumor volume was measured weekly using a digital caliper. LuCaP PDX tumors were validated using STR analysis by Laragen Inc.

### 2.6 Whole Cell Lysate Preparation for Protein Expression

Organoids were grown for 0, 2, and 3 weeks with or without (1 μg/ml) of doxycycline (EMD Millipore, #324385), harvested, and washed twice with cold PBS. Alternatively, LNCaP Crispr knockout cells were harvested using TrypLE and washed twice with cold PBS. The cell pellets were processed in 1% SDS or RIPA/NP40 lysis buffer containing protease and phosphatase inhibitors. After 45 min of suspension at 4℃, pellets were sonicated (Sonication 40-45% amplification, five cycles, 1 s on/off at 4℃). WCL lysate was centrifuged at 10000 rpm for 10mn, and protein estimates were performed using BCA analysis (Pierce Kit: #23225) according to the manufacturer’s instructions.

### 2.7 Immunoblotting

15-30 µg of protein lysate was separated on 4–20% mini-gradient gels and transferred onto polyvinylidene difluoride (PVDF) membranes. Membranes were blocked in 5% milk (Cell Signaling: #9999) in TBST, followed by overnight antibody probing at 4°C °C with RB1 (CST, #9309S), TP53 (Abcam: #PAb240), AR (CST: #5153S), LDH (Novus: #NBP1-48336), KLK3 (CST: #5365), and Histone-H3 (CST: #4499S). All primary antibodies were used at 1:1000 dilution in 5% BSA made in 1X Tris Buffered Saline with Tween (TBST), while secondary antibodies were procured from Thermo Scientific and used at 1:5000 dilution in 1% BSA made in 1X TBST. After primary washing, blots were re-probed with HRP labelled secondary antibodies for an hour at 25℃. The blots were washed with TBST three times for 5 min each and developed using Bio-Rad (Chemiluminescence system).

### 2.8 LDH activity assay

LDH activity was measured using an LDH assay kit (Bio Vision, # K726-500), according to the manufacturer’s instructions. Cell lysates were prepared in the buffer provided with the kit, and equal amounts of protein samples were diluted in 50 μl of the buffer and incubated at room temperature for 10 min. The lysates were further mixed with a reaction mixture (50 μl reaction mix), containing 48μl of development buffer and 2 μl of development enzyme mix and dispensed into a 96-well microplate. OD was measured at 450 nm from both samples and standard with the help of an ELISA plate reader (TECAN pro200, Männedorf, Zürich, Switzerland) at room temperature. Results are plotted as fold-change over the control.

### 2.9 Organoids quantification assays

Organoids (2.5 × 10^3^ cells/well) were seeded in wells (technical replicates) in 3D on 96-well plates and cultured for the indicated number of days in the presence or absence of doxycycline-supplemented media. Quantification was performed using Renilla glow (Promega: #G9681) according to the manufacturer’s protocol.

### 2.10 Seahorse Assay

The extracellular acidification rate (ECAR) and cellular oxygen consumption rate (OCR) were measured using a Seahorse XF96e bioanalyzer (Seahorse Bioscience) and a Glycolytic Rate Assay Kit (Agilent Technologies: #17394939A) according to the manufacturer’s instructions. Organoids and LNCaP Crispr KOs (2.5*10^2^ and 10^4^ cells/well, respectively) were seeded in wells (technical replicates) in a Seahorse XF 96-well assay plate in full growth medium. Post 3^rd^ day, the organoids were supplemented with growth medium containing doxycycline (1 µg/ml). The medium was replaced twice a week for 21 days. Alternatively, 48 h after seeding LNCaP monoclonal Crispr KOs, the medium was carefully washed and replaced with prewarmed sea horse running medium (consisting of nonbuffered DMEM, Agilent Technologies) supplemented with 1 mmol/L sodium pyruvate, 2 mmol/L glutamine, and 10 mmol/L glucose, pH 7.4). Plates were incubated for 60 min in a non-CO_2_ incubator at 37°C before 12 basal measurements were performed. Rot/AA (0.5 mmol/L, inhibitors of the mitochondrial electron transport chain) was immediately added, followed by three additional measurements: 2-deoxy-D-glucose (2-DG), a glucose analog that inhibits glycolysis through competitive binding of glucose hexokinase, was then added with another three further measurements. Post assays, organoid or cell numbers were calculated from each well using Renilla glow or Cyquant assays for RB1-TP53 altered organoid culture or LNCaP monoclonal Crispr KOs. The results from each well were normalized to the respective cell number from the well.

### 2.11 Sample Preparation for Stable isotope resolved metabolomics (SIRM)

Organoids and LNCaP monoclonal Crispr KOs were seeded in 10 cm dishes at 2 million cells per dish and incubated in complete medium. After 3 days, complete media from the organoid culture was supplemented with doxycycline (1μg/ml) for 21 days. The medium was replaced twice a week for 21 days. Post 21-day, organoid cultures were washed with PBS and the medium was replaced with DMEM (Dulbecco’s Modified Eagle’s Medium) containing 15 mM [U-13C]-glucose (Cambridge Isotope laboratories: #CLM-1396-10), 2 mM unlabeled glutamine, and 10% dialyzed FBS (in triplicate), or unlabeled media (15 mM natural abundance glucose; single plate), at 37°C °C in a 5% CO2 atmosphere, RH > 90% for 48 h (approximately one cell doubling). Alternatively, after 48 h, LNCaP monoclonal CRISPR KOs were washed with PBS and the medium was replaced with DMEM (Dulbecco’s Modified Eagle’s Medium) medium containing 10 mM [U-13C]-glucose (Cambridge Isotope laboratories: #CLM-1396-10), 2 mM unlabeled glutamine, and 10% dialyzed FBS (in triplicate), or unlabeled media (single plate), at 37°C °C in a 5% CO_2_ atmosphere, RH > 90% for 24 h (approximately one cell doubling). Glucose concentrations in fresh media were verified by immobilized enzyme electrode amperometry using a YSI 2950 Bioanalyser. After incubation, the medium was aspirated, and the cells (6-8 million per plate) were washed twice with ice-cold PBS and then quenched on a plate with acetonitrile, as previously described. Metabolites were extracted using the Fan Extraction Method [25] with aqueous fractions for NMR and mass spectrometry lyophilized and stored at −80°C until analysis, while the organic fraction was stored at −80°C in chloroform in the presence of butylated hydroxytoluene (BHT). A polar fraction in a 200 µL D_2_O solution containing 0.25 μM DSS-d6 as reference and concentration standard. For each sample, 1D ^1^H Presat and ^1^H-^13^C HSQC spectra were recorded using a phase-sensitive, sensitivity-enhanced pulse sequence using echo/anti echo detection [26] and shaped adiabatic pulses (hsqcetgpsisp2.2) for the ^1^H-^13^C HSQC experiments. The experiments were performed at 293 K with a 16.45 T Bruker Avance Neo spectrometer using a 3 mm inverse triple resonance cryoprobe with an acquisition time of 2 s and a recycle delay of 6 s for the ^1^H experiment, and an acquisition time of 213 ms and recycle delay of 1.75 s for the HSQC experiments. Presaturation was used in each case to suppress the residual water signals. The most concentrated sample from each group was also analyzed by high-resolution 2D multiplicity-edited HSQC-TOCSY and ^1^H/^1^H TOCSY (50 ms mixing time) with an acquisition time in t_2_ of 1 s and a relaxation delay of 1 s and an isotropic mixing time of 50 ms (TOCSY) and an acquisition time of 0.2 s in t_2_, recycle time of 2 s.[27] The nonpolar spectra were acquired similarly except that one half of the nonpolar fraction was evaporated to dryness at RT and re-dissolved in 200 µL of d4-methanol.[28] The nonpolar spectra were acquired similarly, except that half of the nonpolar fraction was evaporated to dryness at RT and re-dissolved in 200 µL of d4-methanol and analyzed as previously described.[25,29]

### 2.12 Mass Spectrometry analysis

ICMS samples were prepared by dissolving the lyophilized powder of the polar fractions in 50 μL of 18 MΩ water. The metabolites were separated with a Dionex ICS6000 system using modifications to the method of Sun *et al* [28] here the eluent was 18 MΩ water at 0.38mL/min, and the KOH gradient was generated electrolytically, then removed by the suppressor post-column. The separated metabolites were analyzed using an Orbitrap Fusion Lumos mass spectrometer (Thermo Scientific). Acetonitrile (0.1 mL) was used as the makeup solvent to assist sample desolvation in the H-ESI (Heated Electrospray Ionization (H-ESI) source. The MS settings were: scan range(m/z) 70-800, resolution 500,000 at m/z 200, RF Lens (%) 30, sheath gas flow rate 50, auxiliary gas flow rate 10, sweep gas flow rate 1, ion transfer tube temperature at 325 °C, vaporizer temperature 350 °C, negative ion voltage at 2500V.

Our in-house metabolite database was incorporated into TraceFinder software (Thermo Fisher) for peak assignments and peak area measurements. A total of 112 metabolite standards were prepared in-house and run at different concentrations to establish calibration curves for each metabolite. Following correction for ^13^C natural abundance (PREMISE software kindly provided by Dr. Richard Higashi, University of Kentucky) and split ratio of the aliquot used for analysis, the calculated amount of each metabolite was then further normalized to the protein quantity measured by BCA assay.

### 2.13 ^18^F-FDG-PET scan

Tumor-bearing mice described under protocol (MIP-006-3-A) were injected with 100 mCi of ^18^F-FDG^+^ in PBS via a tail vein under anesthesia. Sixty minutes after ^18^F-FDG^+^ administration, a whole-body emission PET scan was conducted using BioPET (Bioscan Inc.) under anesthesia with 1.5% isoflurane, with a nominal resolution of 0.375 mm × 0.375 mm × 0.375 mm. Animals were euthanized upon completion of the scan, and tumors and other vital organs were excised from the body, weighed, and biodistribution values were recorded. The total tumor volume and total ^18^FDG^+^ activity were calculated based on the voxel size from the scan. Data were plotted as the total ^18^FDG^+^ activity from each scan, as previously described.[30]

### 2.14 In Vitro Hyperpolarized ^13^C MRS

Hyperpolarized 13C NMR experiments were performed using Spinsolve Benchtop NMR (Magritek). 15-20 million cells from freshly processed PDX tumors were seeded as a suspension culture in a 75 cm ultra-low attachment flask (5 million cells/flask) in 2% Matrigel to form organoids. Alternatively, 10 million LNCaP monoclonal Crispr knockout cells were plated in 15 cm dishes. The cells were incubated in complete medium until the number reached 40-50 million in total. Organoids or cells were collected and washed with PBS. The respective cell numbers were counted and resuspended in 450ul of media. cell suspension was collected in an NMR tube and pre-warmed to 37°C. 75 μL of 96 mmol/L hyperpolarized [1-^13^C] pyruvate solution from a Hypersense DNP Polarizer (Oxford Instruments) were injected into the NMR tube. ^13^C two-dimensional spectroscopic images were acquired using benchtop NMR. Data was plotted as the AUC of hyperpolarized-^13^C-lactate/pyruvate conversion.

### 2.15 In Vivo Hyperpolarized ^13^C MRS/ MRI

*In vivo* hyperpolarized 13C MRI experiments were performed on a 3T MRI Scanner (MR Solutions Inc.) using a 17-mm diameter home-built ^1^H/^13^C coil. A 96 mmol/L hyperpolarized [1-^13^C] pyruvate solution from a Hypersense DNP Polarizer (Oxford Instruments) was administered to a male NODscid gamma (NSG) mouse under anesthesia with 1.5% isoflurane via a tail vein cannula (12 mL/g body weight), as described in the animal protocol (RBB-159-3-P). ^13^C two-dimensional spectroscopic images were acquired with a 32 × 32 mm field of view in an 8-mm axial slice. [30]

### 2.16 Statistical analysis

All experiments were performed with at least three biological replicates per replicate. Data are displayed as the mean ± SEM or mean ± SD as indicated. Statistical significance (p < 0.05) was determined using Student *t* t-test and one-way ANOVA using GraphPad Prism Software, as appropriate. The PC and PLS-DA analyses on z-transformed metabolites were performed using Viime.[31]

## 3. Results

### 3.1 NEPC cannot be conclusively distinguished from adenocarcinoma by ^18^FDG uptake alone in *in vitro* models

To broadly characterize ^18^FDG uptake in an extensive cohort of CRPC organoids relative to prostate cancer phenotype and genotype, we measured ^18^FDG uptake over 120 minutes in PDX-derived ARPC and NEPC organoids (n=18 cell lines), patient biopsy-derived ARPC organoids (n=2), amphicrine (a CRPC subtype that maintains AR activity while also expressing classic NE markers,[32,33] n=1) and ARPC cell lines (n=3). Figure 1A shows the percentage of ^18^FDG uptake normalized to the number of live cells for each model. Although the final ^18^FDG uptake values showed variation among the biological replicates, we observed at the final time point a clear separation between models with high and low ^18^FDG uptake, with a separation line occurring around 6% uptake (see Figs. 1A and S1A). While the distinction between high and low ^18^FDG uptake groups is evident, the correlation between ^18^FDG uptake and phenotype is not as straightforward. On average, adenocarcinoma (ARPC) models exhibited higher ^18^FDG uptake compared to NEPC models (8.1 ± 1.5% vs. 4.7 ± 2.1% SEM). With the exception of the LuCaP 145.1 organoids, all NEPC models fell into the low ^18^FDG uptake group. By contrast, adenocarcinoma models were almost equally distributed between the high and low ^18^FDG uptake groups. Therefore, it is not possible to definitively distinguish lineage in organoid cultures based solely on ^18^FDG uptake.

**Figure 1.**
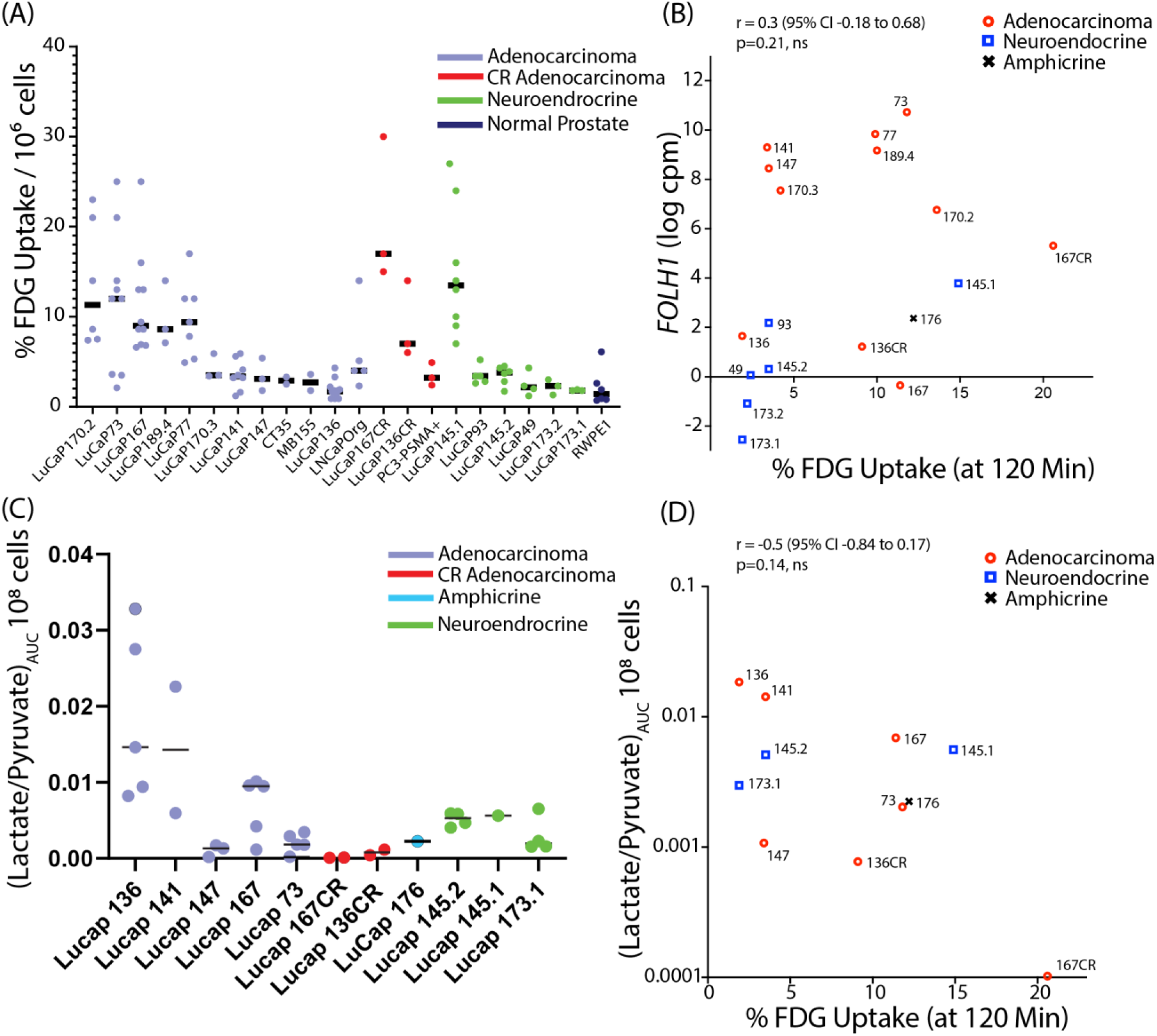
^18^FDG uptake does not correlate with hyperpolarized ^13^C lactate/pyruvate conversion. **A.** Percent ^18^FDG uptake at the final timepoint of 120 minutes for biological replicates of the same model. **B.** Pearson correlation of ^18^FDG uptake at the final timepoint (120 minutes) against *FOLH1* transcription (cpm) in LuCaP organoids **C.** AUC of hyperpolarized-^13^C-lactate/pyruvate conversion in LuCaP organoids. **D.** Scatterplot of ^18^FDG uptake at the final timepoint (120 minutes) against the AUC of hyperpolarized-^13^C-lactate/pyruvate conversion in LuCaP organoids.

The complex genomic variability among clinical ARPC and PDX ARPC models complicates comparisons between castration-resistant and castration-sensitive models.[22] The extent of metabolic reprogramming during disease progression can be determined more directly by investigating longitudinal changes in experimentally castrated PDX-derived models. In line with reported clinical findings (*15*), the castration-resistant (CR) models LuCaPs136CR and 167CR showed higher retention of ^18^FDG compared to their castration-sensitive (CS) counterparts, LuCaPs 136 and 167 (Figs. 1D, S1B and C). In contrast, castration-sensitive models displayed relatively high pyruvate-to-lactate conversion compared to their castration-resistant counterparts in the limited direct comparisons that were available.

### 3.2 PSMA protein levels do not correlate with prostate cancer phenotype or ^18^FDG uptake in *in vitro* models

^18^FDG imaging is not the only molecular imaging technique used for prostate cancer staging. PSMA targeted imaging is increasingly used in the clinic, thus, we compared *FOLH1* RNA levels, a reasonable surrogate for PSMA protein expression in LuCaP models,[14] to ^18^FDG uptake in each model (Fig. 1B). As with ^18^FDG, there was a clear separation between high and low *FOLH1* groups. As expected from previous preclinical studies, ARPC models generally showed high *FOLH1* levels and *RB1* null models, including NEPC models, demonstrated markedly lower levels of FDG uptake, although there were two exceptions, NEPC LuCaP145.1 and ARPC LuCaP167. No correlation was found between ^18^FDG uptake and *FOLH1,* which matches the discordance between ^18^FDG-PET and PSMA imaging observed in the clinic.[34]

### 3.3 Glucose Uptake and Aerobic Glycolysis Are Not Linked in Prostate Cancer Organoid Models

^18^FDG uptake is restricted to assessing glucose uptake and phosphorylation and lacks sensitivity for other, more downstream metabolic processes.[35] In contrast, hyperpolarized MRI specifically measures the forward flux of pyruvate conversion to lactate through lactate dehydrogenase, providing a more direct means of evaluating a glycolytic phenotype than ^18^FDG uptake (Fig 1C). Similar to ^18^FDG uptake, there was high variability between samples with a clear division between high and low pyruvate-to-lactate conversion, which did not correlate well with adenocarcinoma/NEPC phenotypes. No correlation was found between ^18^FDG uptake and lactate/pyruvate flux, demonstrating that glucose uptake in these models is not correlated with aerobic glycolysis (Warburg physiology) (Fig. 1D).

### 3.4 AR expression is maintained after *RB1/TP53* knockdown in castration sensitive cell lines and PDX organoids in vitro

The models in Fig. 1 incorporate a wide range of prostate cancer phenotypes in a variety of genetic contexts. Although our initial characterization included both CRPC and castration-sensitive models, the experimental limitations of CRPC lines necessitated our focused genetic investigations in castration-sensitive systems. CRPC lines grow poorly in culture and fail to establish stable organoids for the duration required for detailed *in vivo* experiments, primarily due to genomic instability where organoids frequently lose proliferative capacity beyond early passages despite initially maintaining cancer hallmarks.[22,36–39] This genomic instability and phenotypic drift in CRPC models led us to employ castration-sensitive systems where stable genetic profiles enabled precise interrogation of *RB1/TP53* effects. To more narrowly look at the specific effect of *RB1* and *TP53* on molecular imaging in a traditional cell line model, we used CRISPR/CAS9 knockout of *RB1* and/or *TP53* in castration sensitive LNCaP cells (RB1^+/+^/TP53^+/+^). In this model, both *RB1* and *TP53* expression levels were effectively null (Fig. S2B), confirming it as an efficient model of *RB1* and *TP53* knockdown. To examine the effects of *RB1* and/or *TP3* loss in a more physiologically representative organoid model, we introduced stable short hairpin RNAs (shRNAs) targeting *RB1* and *TP53* into castration sensitive LuCaP 167 PDX-derived organoids (RB1^-/+^/TP53^+/+^).[22,40], which retain stable growth characteristics across passages compared to CRPC-derived organoids. Doxycycline-induced shRNA effectively depleted *RB1* transcription in LuCaP 167, while shRNA induction reduced but did not eliminate the expression of TP53 (Fig. S2A). LuCaP 167 constitutively expresses high levels of the androgen receptor splice variant AR-V7, implying AR dependent disease progression.[41] Expression of AR was retained upon depletion or loss of *RB1* and/or *TP53* compared to the control, while expression of the AR downstream target KLK3 decreased after doxycycline induction of *RB1* (Fig. S2C). This mirrors previous reports in CRPC that while inactivation of *RB1* suppresses AR signaling, AR expression is maintained after *RB1* loss.[18]

### 3.5 *RB1/TP53* knockdown does not affect ^18^FDG uptake in LNCaP models *in vitro* or LuCaP PDX castration sensitive ARPC mouse models

Having established models of RB1 and TP53 knockdown in both traditional cell line and patient-derived organoid models, we explored how these alterations affect glucose uptake. In the conventional LnCaP cell line, CRISPR/CAS9 knockout of *RB1* or *TP53* did not affect ^18^FDG uptake *in vitro* (Fig. S3). To investigate the impact of RB1/TP53 loss in in a more physiologically representative model, we evaluated ^18^FDG PET scans of LuCaP 167 organoid-derived PDX CRPC tumors subjected to *RB1/TP53* knockdown. The PET scans, shown in Fig.2, revealed substantial FDG uptake in the CRPC tumors, with no significant differences between the RB1/TP53-depleted and control groups. To obtain a more precise measurement of FDG uptake, we excised these tumors after the scan and measured tissue uptake with a gamma counter. The results confirmed that there was no statistically significant difference in FDG activity after knockdown of either *RB1, TP53,* or *TP53* and *RB1* (Fig. S4). *RB1/TP53* knockdown did not significantly affect ^18^FDG uptake in either prostate cancer model tested.

**Figure 2.**
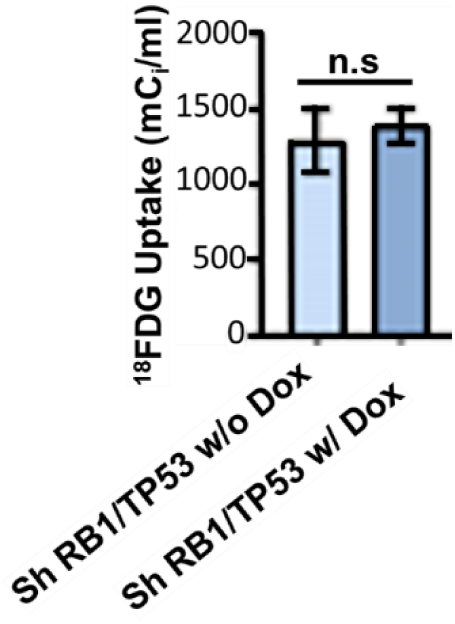
Combined *RB1*/*TP53* knockdown does not affect ^18^FDG uptake in prostate cancer mice xenografts. Total ^18^FDG activity in LuCaP 167 organoids PDX mouse xenograft models with (n=7) or without (n=3) induction of shRNA targeting *RB1* and *TP53* by doxycycline. Data mean ± SEM, ns = not significant. Student t test was performed using Prism 9.0. p= * (0.05), ** (0.01), *** (0.001).

### 3.6 Basal respiration and glycolytic activity are increased following decreased RB1 expression

As *RB1* loss is often associated with increased glycolysis, we used Seahorse assays to characterize the metabolic phenotype of *RB1* and/or *TP53* altered cell lines and organoids. Basal and maximal respiration were significantly elevated for both *RB1* and *RB1/TP53* alterations compared to the control for both LuCaP167 organoids and LnCaP cells (Fig. 3A and C). The basal and maximal respiration of the combined *RB1/TP53* alterations was significantly higher than *RB1* only, while partial depletion of *TP53* in LuCaP167 or complete loss of *TP53* in LNCaP resulted in no change. Extracellular acidification (ECAR), a measure of proton efflux from both glycolysis and respiration and a surrogate for lactate export, was also higher in *RB1* and *RB1/TP53* altered models (Fig. 3B and D).

**Figure 3.**
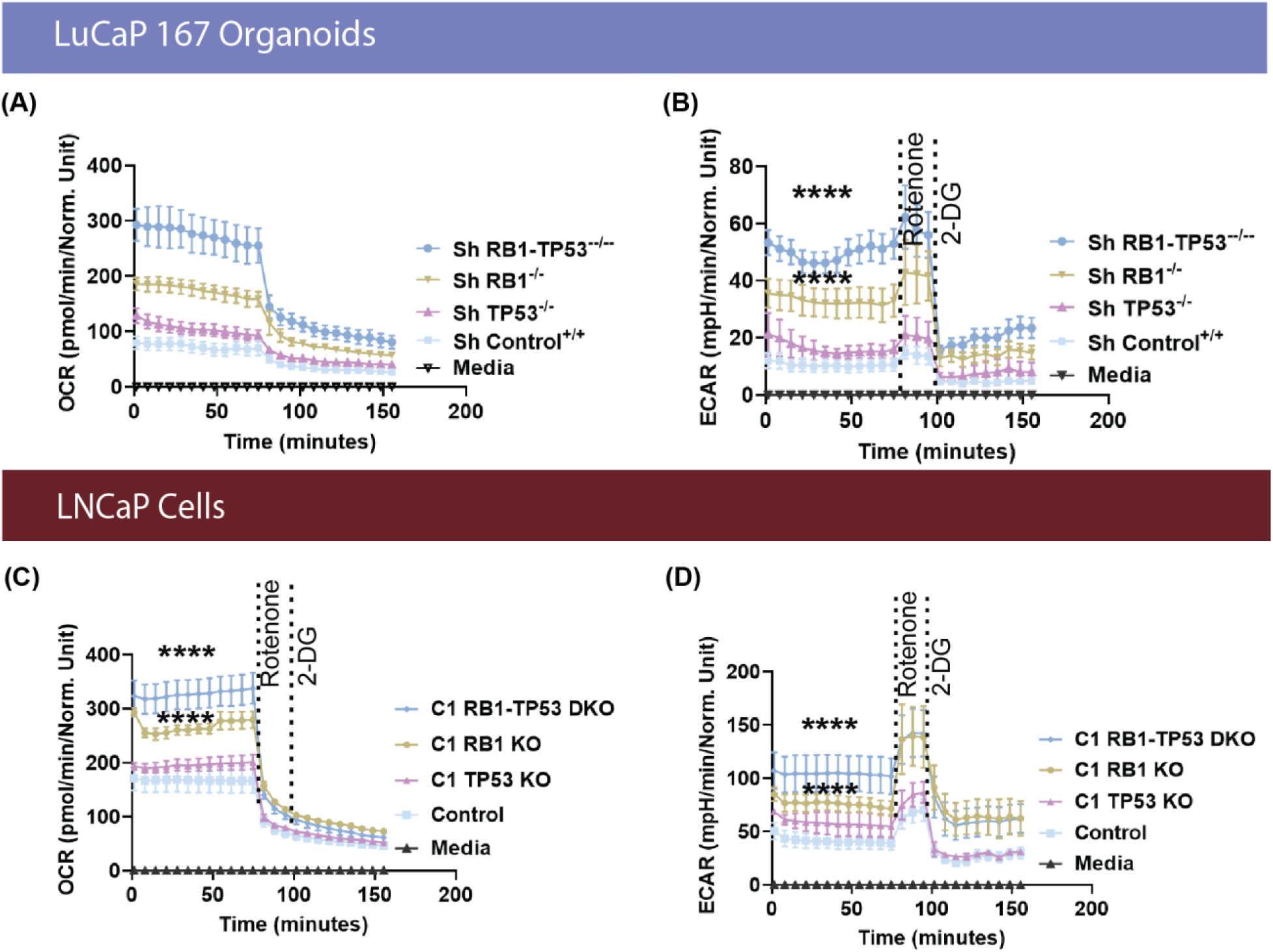
Depletion of *RB1* alone or combined with *TP53* elevates basal respiration rates and increased glycolytic activity. (**A)** Oxygen consumption rate (OCR) and (**B)** extracellular acidification rate (ECAR) of LuCaP167 organoid cultures either with 3 weeks of doxycycyline induction of shRNA targeting *RB1* and / or *TP53* or without (control). using a Glycolytic Rate assay kit on a Seahorse XF96e analyzer. **(C and D)** OCAR and ECAR performed with *RB1* and/ or *TP53* genetically-modified LNCaP Crispr KOs. Data are represented as mean ± SEM, n=3 expr. ns = not significant. One-way ANOVA test with Dunnett’s multiple comparisons test was performed using Prism 9.0. p= * (0.05), ** (0.01), *** (0.001).

The large increase in the extracellular acidification rate in the combined *RB1/TP53* modified models is characteristic of an increased glycolytic phenotype, as meeting a steady-state ATP demand exclusively by glycolysis is considerably more acidifying than meeting the same demand strictly by oxidative phosphorylation. To further analyze the mechanism of increased acidification, we evaluated the enzymatic activity of lactate dehydrogenase (LDH) (Fig. 4), an enzyme that converts pyruvate to lactate. Consistent with the Seahorse assay results, increased LDH activity was observed upon depletion or loss of *RB1* in both models. Overall, the data is consistent with the loss of *RB1* with and without *TP53* loss leading to increased metabolic activity.

**Figure 4.**
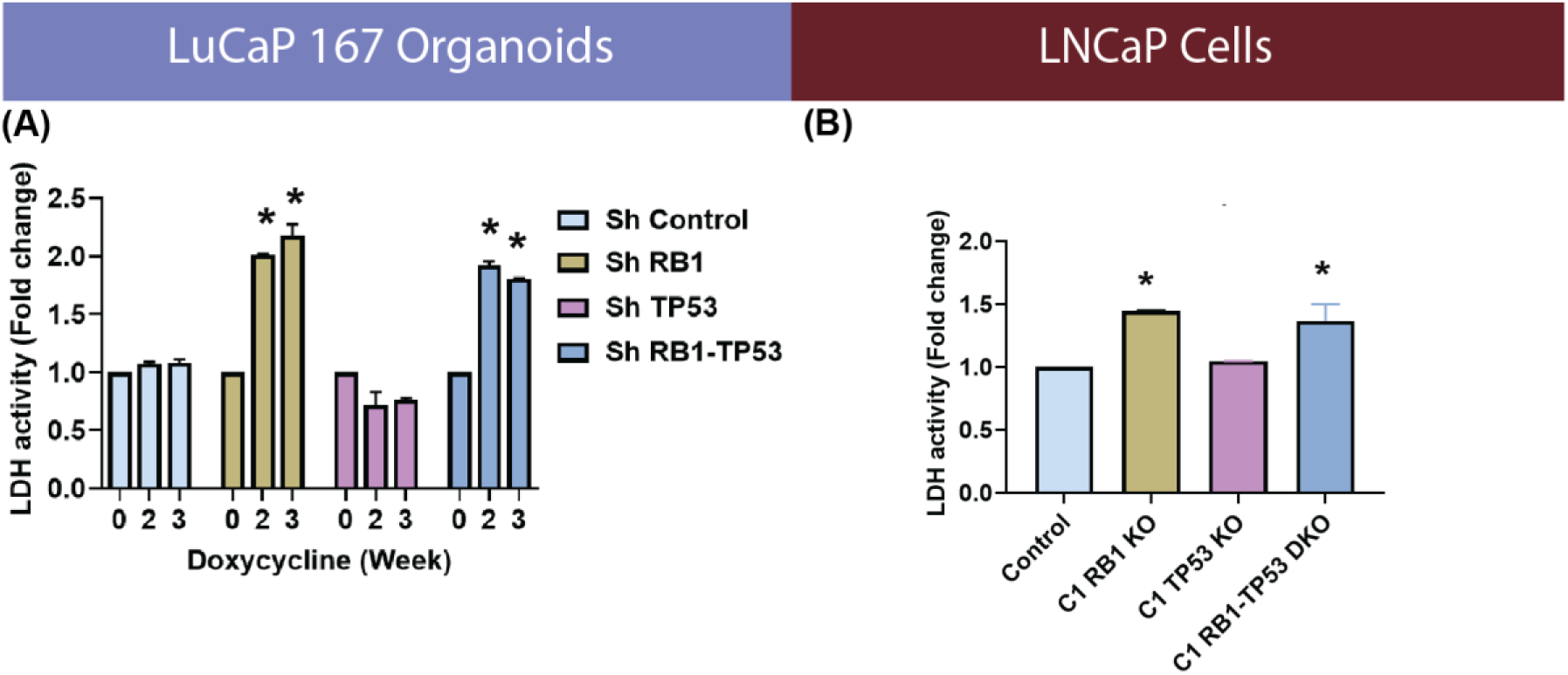
RB1 depletion leads to increased LDH activity. ELISA based LDH activity of **(A)** LuCaP167 organoids with or without ShRNA targeting *RB1* and/or *TP53*, and **(B)** LNCaP Crispr-mediated knockout (KO) of *RB1* and/or *TP53*. One-way ANOVA test with Dunnett’s multiple comparisons test was performed using Prism 9.0. p= * (0.05), ** (0.01), *** (0.001).

### 3.7 Combined *RB1/TP53* depletion leads to metabolic changes, including a diversion of glucose into glycogenesis

To gain a more comprehensive understanding of the metabolic transformation after *RB1* and/or *TP53* depletion, we quantified the downstream metabolites of ^13^C glucose in the LuCaP 167 organoid model by NMR (Fig S5). ^13^C glycogen was highly enriched after dual *RB1/TP53* depletion while steady state concentrations of ^13^C glucose decreased, suggesting a diversion of glucose to form glycogen. It also is of interest that glutamine levels increased since glutamine utilization has been shown to suppress glucose uptake in various cancer models.[42] ^13^C UDP-glucose, an important precursor in the glycogenesis pathway, was found in high abundance but was unaffected by either *RB1* or *TP53* depletion. Consistent with the Seahorse and LDH activity assays, ^13^C labeled lactate concentrations were elevated after dual *RB1/TP53* depletion with more modest increases occurring following single gene modifications. In the non-polar fraction, both de novo cholesterol synthesis and ^13^C incorporation into lipid acyl chains and glycerol headgroups were significantly elevated after depletion of either *RB1* or *TP53*, which is noteworthy since cholesterol is a precursor for androgen synthesis[43] and indirectly influences many signaling pathways through the formation of raft domains.[44] Increased de novo fatty acid synthesis is also a hallmark of prostate cancer metabolism and is correlated with progression and metastasis,[45] although it is uncertain whether it is a driving factor or an epiphenomenon related to increased cholesterol production, which shares some of the same synthetic pathways.[46]

When measured by ^13^C NMR, glucose metabolism in the LnCaP model was less affected by *RB1/TP53* loss relative to LuCaP167 (Fig. S6). Of the 12 metabolites whose ^13^C incorporation was high enough to be quantified by ^13^C HSQC 1D NMR, only glycogen and cholesterol showed significant (roughly 3-fold) differences after the loss of both *RB1* and *TP53*. This is not evidence that other changes did not occur, as ^13^C NMR has limited sensitivity and can only detect metabolites with high steady-state concentrations. Therefore, we repeated the ^13^C glucose tracer experiment using targeted IC-MS to detect changes in intermediates in the glycolytic and TCA pathways, the low abundance of which is problematic for NMR.

In the LuCaP 167 organoid models, *RB1* and *TP53* depletion was clearly differentiated from control samples by IC-MS (Figs. 5A and S7), primarily by a decrease in pyruvate and an increase in lactate concentrations. Intermediates in the latter half of the TCA cycle decrease upon depletion of either *RB1* or *TP53*. Glucose-1-Phosphate, one of an intermediate of glycogenesis, robustly increased following *RB1* depletion. Thus, both ^13^C HSQC 1D NMR and IC-MS detected many of the same metabolite concentration shifts, as observed with ^13^C NMR.

**Figure 5.**
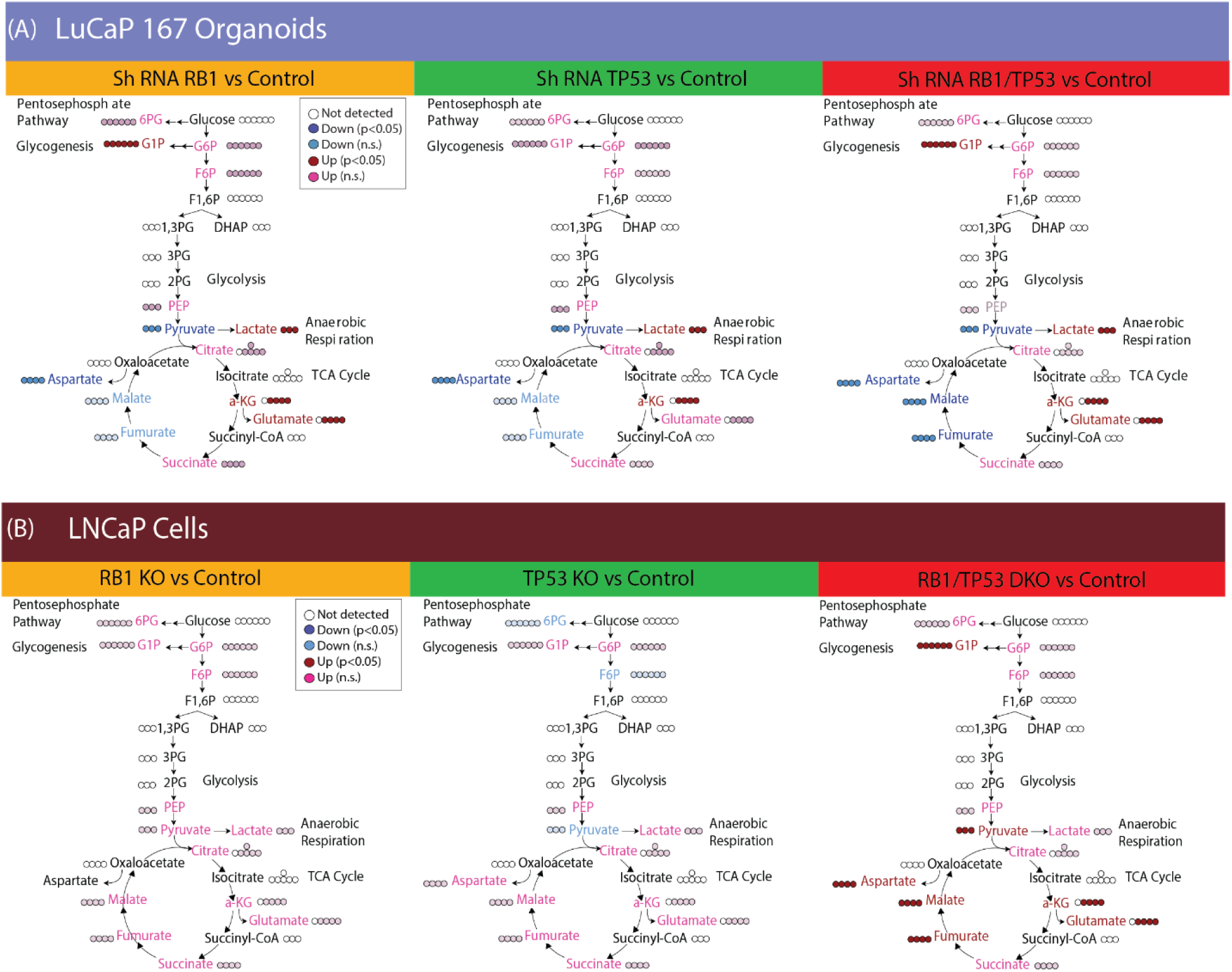
Metabolic changes upon *RB1* and/ or *TP53* depletion by ICMS. **A.** LuCaP167 organoids either with or without ShRNA targeting RB1 and/or TP53 **(B)** LNCaP Crispr-mediated knockout (KO) of RB1 and/or TP53.

The primary difference in metabolic reprogramming for the LnCaP cells was observed in the combined *RB1/TP53* null samples (Figs. 5B and S8). Dual *RB1/TP53* knockout increased activity throughout the TCA cycle, which was not seen to a significant extent with individual knockouts. Consistent with the ^18^FDG uptake data, levels of glucose-6-phosphate were unchanged following *RB1* and/or *TP53* alterations. The glycogenic intermediate, glucose-1-phosphate increased, as expected from the ^13^C HSQC 1D NMR results.

To summarize, *RB1* loss in different castration sensitive prostate cancer models led to consistent changes including: 1) increased lactate, reflecting higher lactate dehydrogenase activity, 2) increased glucose-1-phosphate and glycogen levels, suggesting redirection of glucose metabolism to gluconeogenesis, and 3) modulation of the TCA cycle, specifically a consistent increase in αKG and glutamate across models in addition to other model-specific changes.

### 3.8 Increase in LDH flux after *RB1* knockdown can be detected in vivo by ^13^C-HPMRS

NMR and ICMS identified several metabolites that were upregulated after *RB1/TP53* alteration. To determine whether pyruvate-to-lactate flux imaging could be a tractable approach for analyzing such a phenotype *in vivo*, we first assayed hyperpolarized ^13^C-pyruvate conversion in genetically modified LnCaP cells in vitro (Fig. S9A). We observed increased pyruvate flux to lactate in *RB1* null cells and an additional increase in combined *RB1/TP53* null cells, but no change in *TP53* null only cells. To extend this data, we assayed LuCaP167 organoid-derived PDX tumors *in vivo* (Fig. 6) using magnetic resonance spectroscopy to detect the de novo generation of new metabolites from pyruvate. In LuCaP 167 tumors, the rate of pyruvate to lactate conversion reflective of LDH flux was significantly increased with depletion of *RB1*(see Fig. S9B) with or without partial depletion of *TP53*. We conclude that measuring lactate flux may be marker of *RB1* modified tumors, which are known to be associated with rapid progression.

**Figure 6.**
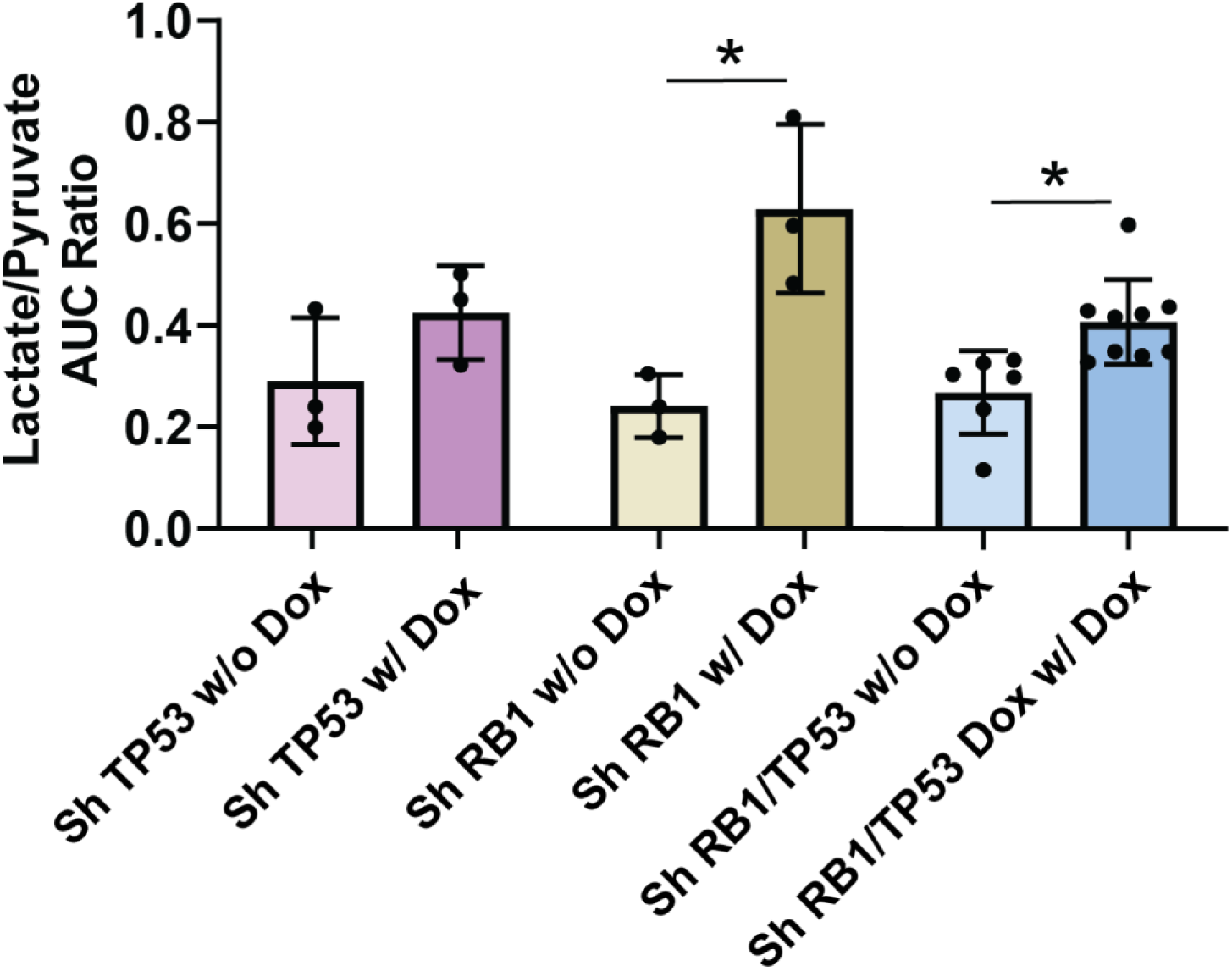
^13^C-HPMRS shows depletion of *RB1* alone or combined with *TP53* depletion increases LDH flux in vivo. *In vivo* LDH activity in LuCaP 167 organoids PDX mouse xenografts with or without doxycycline-inducible shRNA knockdown of *RB1* and/or *TP53* calculated from time-resolved ^13^C hyperpolarized non-localized spectra from the ratio of the area under the lactate and pyruvate signals. Data are represented as mean ± SD, ns = not significant Student t test was performed using Prism 9.0. p= * (0.05), ** (0.01), *** (0.001).

## 4. Discussion

CRPC is a highly heterogeneous tumor type with variability not only in the underlying genomic landscape, but also in the lineage phenotype.[12,13] Although tumor imaging modalities can be powerful methods to both detect tumors and gain additional predictive or prognostic information, it is important that such findings be validated at a molecular level. Here we take advantage of a large cohort of tractable, reproducible organoid models to evaluate for the first time the distribution of glycolytic phenotypes among multiple phenotypically and genetically characterized models.[22,47] using assays that quantify the initial and final steps of glycolysis, FDG uptake, and pyruvate-to-lactate conversion, respectively. Although the highest levels of FDG uptake were observed in ARPC, while uptake in NEPC models was generally minimal, we found that FDG cannot distinguish between ARPC or NEPC lineage phenotypes *in vitro* due to substantially overlapping ranges in values. This is consistent with the previously described role of AR in stimulating the expression of glycolytic proteins.[4,5] Of particular interest, there was a clear lack of correlation between glucose import and lactate production, indicating that FDG uptake is not necessarily related to aerobic glycolysis (Warburg physiology) (Fig. 1D). Experimental models have shown that lactate production and glucose import are modulated in parallel directions in response to genetic manipulation.[48] However, it is perhaps not unexpected that intermediary metabolites connected by multiple pathways have differences in rate-limiting steps among patient-derived samples.

PSMA-PET is a widely used marker of prostate cancer and is generally used to identify ARPC and CRPC. It is often used in combination with FDG-PET, as PSMA uptake can be reduced or completely suppressed in the later stages of CRPC, while FDG-PET of the same lesions is usually positive in advanced PSMA-negative prostate cancer. Since PSMA PET is a biomarker for the effectiveness of PSMA-directed therapies, i.e., PSMA-targeted radioligand therapy, PSMA-/FDG+ discordant cases tend to predict poorer outcomes, [49,50] although the importance of this discordance has recently been questioned.[51] Independent of PSMA PET, FDG PET uptake is also associated with poorer prognosis in prostate cancer, although the cellular mechanisms underlying the increased glucose metabolism remain poorly understood.[52]

Given the prognostic significance of FDG uptake, it is important to explore the genetic factors that may influence this metabolic characteristic. Functional *RB1* loss in CRPC is the most significant predictive mutation with respect to both poor survival and lack of AR signaling inhibitor drug responsiveness for adenocarcinoma pathologies.[14,53] Combined *RB1/TP53* loss is an almost universal feature of lethal NEPC. [15,16] To address whether the loss of *RB1* and/or *TP53* in adenocarcinoma leads to increased FDG uptake and/or glycolysis, we analyzed two castration sensitive AR+ adenocarcinoma models that were genetically depleted for these molecules. While CRPC models would more closely mirror advanced disease, the genomic instability and phenotypic drift in CRPC models directed our approach toward castration-sensitive systems, which provide consistent baseline metabolic signatures essential for reliable interpretation of genetic manipulation effects on glucose metabolism.

We determined that *RB1* loss in castration sensitive models was associated with increased extracellular acidification, a measure of glycolytic lactate production, and increased oxygen consumption, a measure of oxidative phosphorylation (Figure 3). Consistent with this, LDHA enzyme activity (Figure 4) was increased, as was cellular lactate itself (Figure 5), as determined by ICMS. Increased expression of LDHA and MCT4, the lactate efflux transporter, have been described for prostate cancer progression in experimental models and clinical samples.[54,55] Other properties that were observed in both models included increased glucose diversion into glycogen synthesis and increased membrane lipid synthesis. We assessed whether quantitative changes to FDG or HP-MRI imaging were correlated with the genomically-driven metabolic alterations in glycolytic phenotype. Consistent with increased lactate production, ^13^C HP-MRI spectroscopy signals were increased following *RB1/TP53* depletion. On the contrary, FDG uptake values remained the same. This data suggests that RB1 expression limits LDHA-dependent lactate production but does not affect glucose availability or uptake.

Clinical experience has shown that increased FDG uptake in CRPC patients is associated with poor survival. We observed a poor correlation between FOLH1 expression and FDG uptake, indicating that PSMA scans cannot be used to predict FDG uptake. Furthermore, we show here that cancer cells themselves do not demonstrate increased FDG uptake as in *RB1/TP53* null NEPC organoids derived from patients or ARPC organoids in which *RB1/TP53* was depleted. How does one explain the observation that clinical FDG uptake is increased in neuroendocrine and other advanced prostate cancer phenotypes, with our finding that FDG uptake was unchanged despite genetic alterations associated with poor survival in ARPC and a high rate of transition to NEPC? One potential explanation relates to the limitations inherent in our organoid models. While valuable, they may not fully capture the metabolic programming aspects of NEPC seen in vivo. Our findings demonstrate two key points: 1) there was no clear relationship between lineage (adenocarcinoma vs. neuroendocrine) and FDG uptake in our models (Figure 1), and 2) in an AR-positive, castration-sensitive context, *RB1/TP53* loss did not increase FDG uptake (Figure 2 and S3). Although *RB1/TP53* loss is almost universally observed in the transition to NEPC, our results suggest that this genetic alteration alone is insufficient to drive the increased FDG uptake observed clinically. This discrepancy might arise because the metabolic effect of *RB1/TP53* loss is context-dependent, requiring factors absent in our models such as castration resistance, altered AR signaling, or synergy with other genetic drivers (e.g., *MYCN* amplification or *PTEN* loss). Alternatively, the high clinical FDG uptake could be driven by entirely different regulatory circuits controlling glucose metabolism, activated independently of *RB1/TP53* status.

Another significant factor contributing to clinical FDG uptake, and perhaps a more dominant effect in vivo, involves the tumor microenvironment (TME). FDG uptake has been shown to be highly dependent on microenvironmental host immune cells such as myeloid-derived macrophages and activated T cells.[42] These cells are glucose-avid and tend to outcompete tumor cells for glucose uptake in the context of the tumor microenvironment. Importantly, acidic tumor microenvironments secondary to lactate accumulation promote M2 macrophage and Treg differentiation, while suppressing T effector cell function, leading to an immunosuppressive microenvironment.[56] These activated TME cells are metabolically active and likely take up FDG avidly. It will be of interest in the future to determine in clinical samples whether there are correlations between 1) advanced prostate tumors with decreased AR signaling and/or 2) *RB1* loss of function, acidic tumor microenvironments, increased M2 macrophage numbers, and associated increased FDG uptake.

## 5. Conclusions

While *RB1/TP53* loss drives significant metabolic reprogramming in prostate cancer - including increased glycolysis, elevated lactate production, enhanced TCA cycle activity, and glucose diversion into glycogen synthesis - these extensive metabolic alterations are not reflected in conventional ^18^FDG-PET imaging uptake values. This metabolic plasticity highlights the importance of considering multiple metabolic pathways when developing therapeutic strategies for prostate cancer, particularly in tumors with *RB1/TP53* alterations. This suggests multimodal molecular imaging approaches may provide more accurate characterization of *RB1*-deficient tumors and better guide treatment decisions for CRPC patients.

---

**Supplementary Figure 1.**
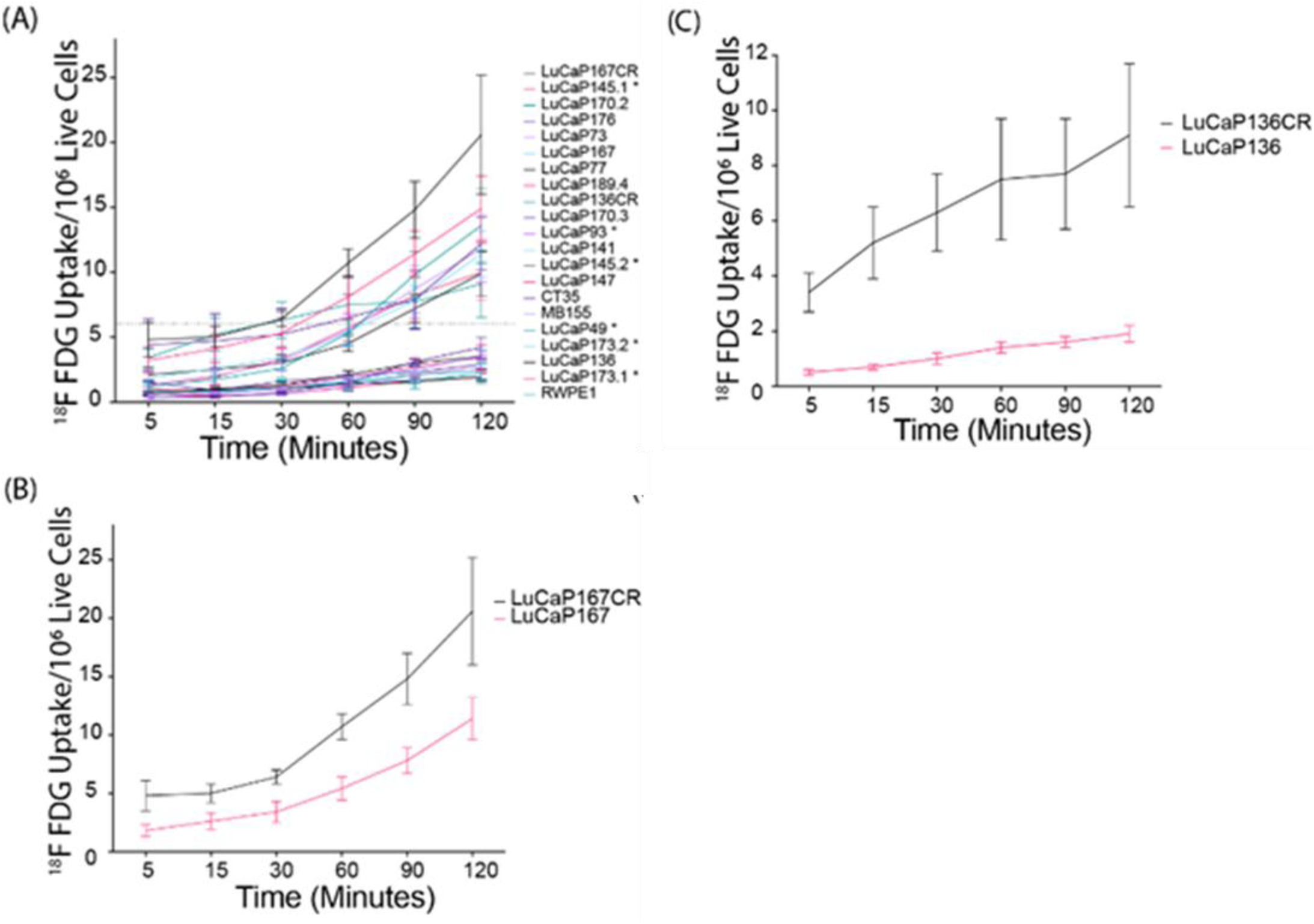
Adenocarcinoma, castration-resistant prostate cancer models display higher FDG uptake. **(A)** FDG uptake over time per model; % uptake / 10^6^ live cells = [(cpm uptake) / (cpm total added) x (live cells total added)] x (10^6^live cells) x 100) for each timepoint. Line denoting the 6% FDG uptake indicates the separation between FDG-high models and FDG-low models. **(B, C)** FDG uptake over time in matched parental (castration-sensitive) and castration-resistant (CR) PDX-derived organoid models, where CR was selected via *in vivo* androgen deprivation (surgical castration of host mice).

**Supplementary Figure 2.**
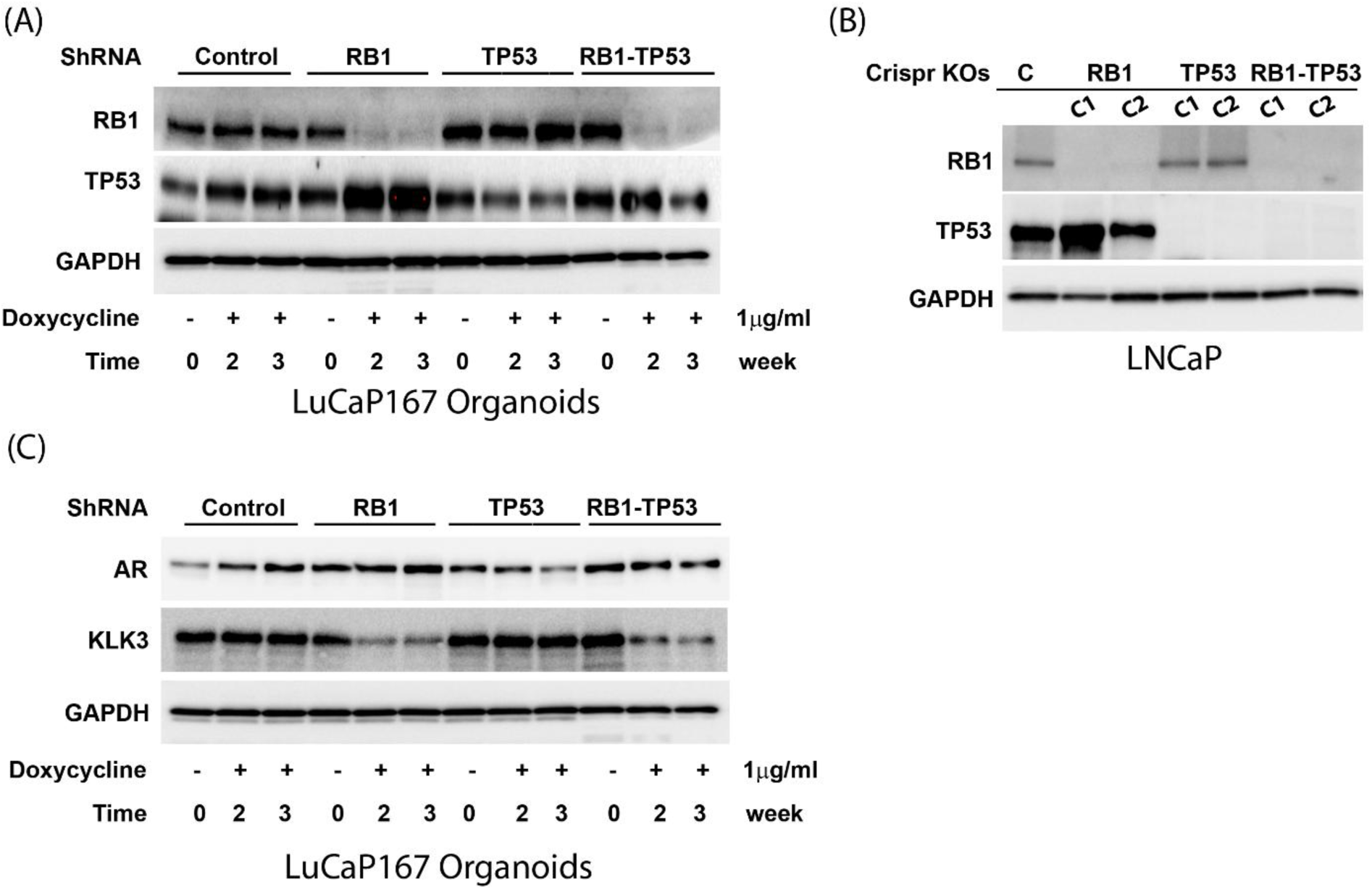
*RB1* and/ or *TP53* depletion maintains AR expression. (**A)** Western blot showing RB1, TP53 and GAPDH protein expression in *RB1* and/or *TP53* altered LuCaP167 organoid cultures after 0, 2, or 3 weeks of ShRNA induction by doxycycline. **(B)** Western blot showing RB1, TP53 and GAPDH protein expression in *RB1* and/or *TP53* LNCaP Crispr KOs. **(C)** Western blot showing AR, KLK3 and GAPDH protein expression in *RB1* and/or *TP53* altered 167 LuCaP organoid cultures post 0, 2, or 3 weeks of doxycycline induction.

**Supplementary Figure 3.**
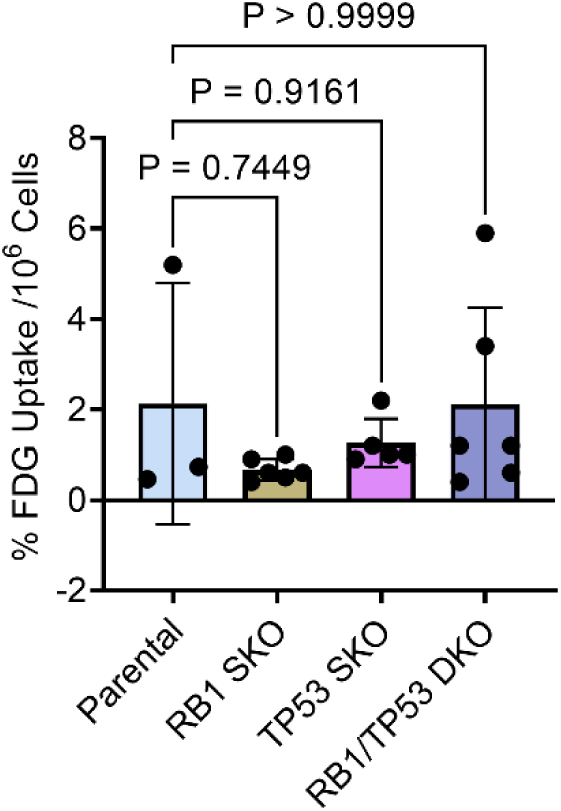
*RB1* and/ or*TP53* knockdown does not affect: ^18^FDG^+^ uptake in LnCaP monoclonal Crispr knockouts. ^18^FDG uptake per million cells after 120 minutes CRISPR/CAS9 knockout of *RB1* and/or *TP53*. Data is mean ± SD. One-way ANOVA test with Dunnett’s multiple comparisons test was performed using Prism 9.0. p= * (0.05), ** (0.01), *** (0.001).

**Supplementary Figure 4.**
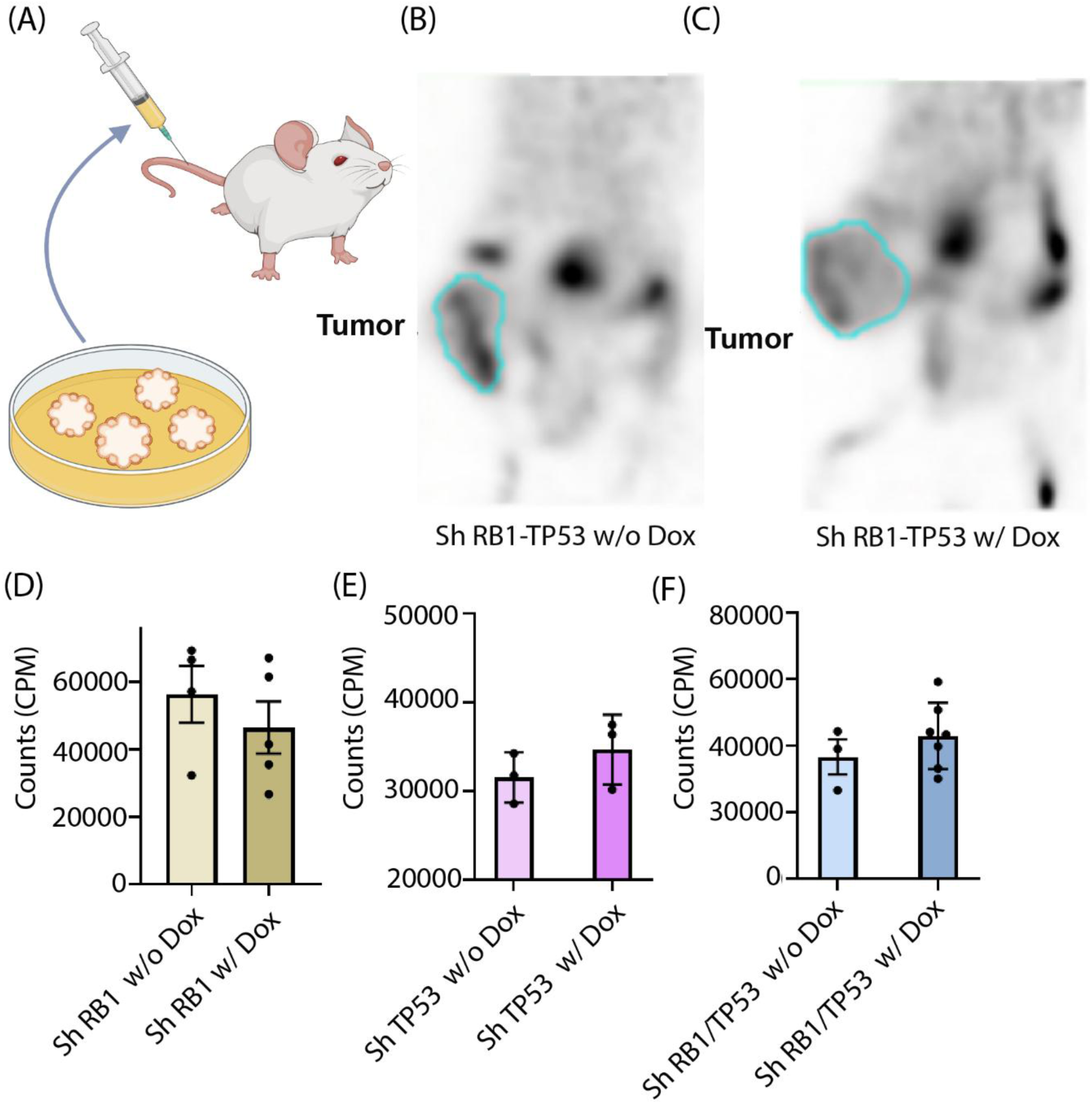
*RB1* and/ or*TP53* loss does not affect ^18^FDG^+^ uptake in a LuCaP 167 PDX model. **(A)** Experimental setup. LuCaP 167 organoids were cultured in vitro, mixed with Matrigel, and implanted subcutaneously into NOD scid gamma (NSG) mice to create tumor xenografts. **(B and C)** ^18^FDG^+^-PET scan image showing total ^18^FDG^+^ activity present inside LuCaP 167 (PDX) mouse castration sensitive prostate cancer xenograft models with and without induction of shRNA targeting *RB1* and *TP53* by doxycycline. **(D, E, F)** Total amount of ^18^FDG^+^ calculated in scintillation counter from excised tumors with shRNA targeting *RB1* (n=5), *TP53* (n=3), *RB1/TP53* (n=7) induced by doxycycline, and their respective controls without doxycycline induction (n=4, 3, 3). Difference between control and knockdowns were not statistically significant. Data mean ± SEM, ns = not significant. Student t test was performed using Prism 9.0. p= * (0.05), ** (0.01), *** (0.001).

**Supplementary Figure 5.**
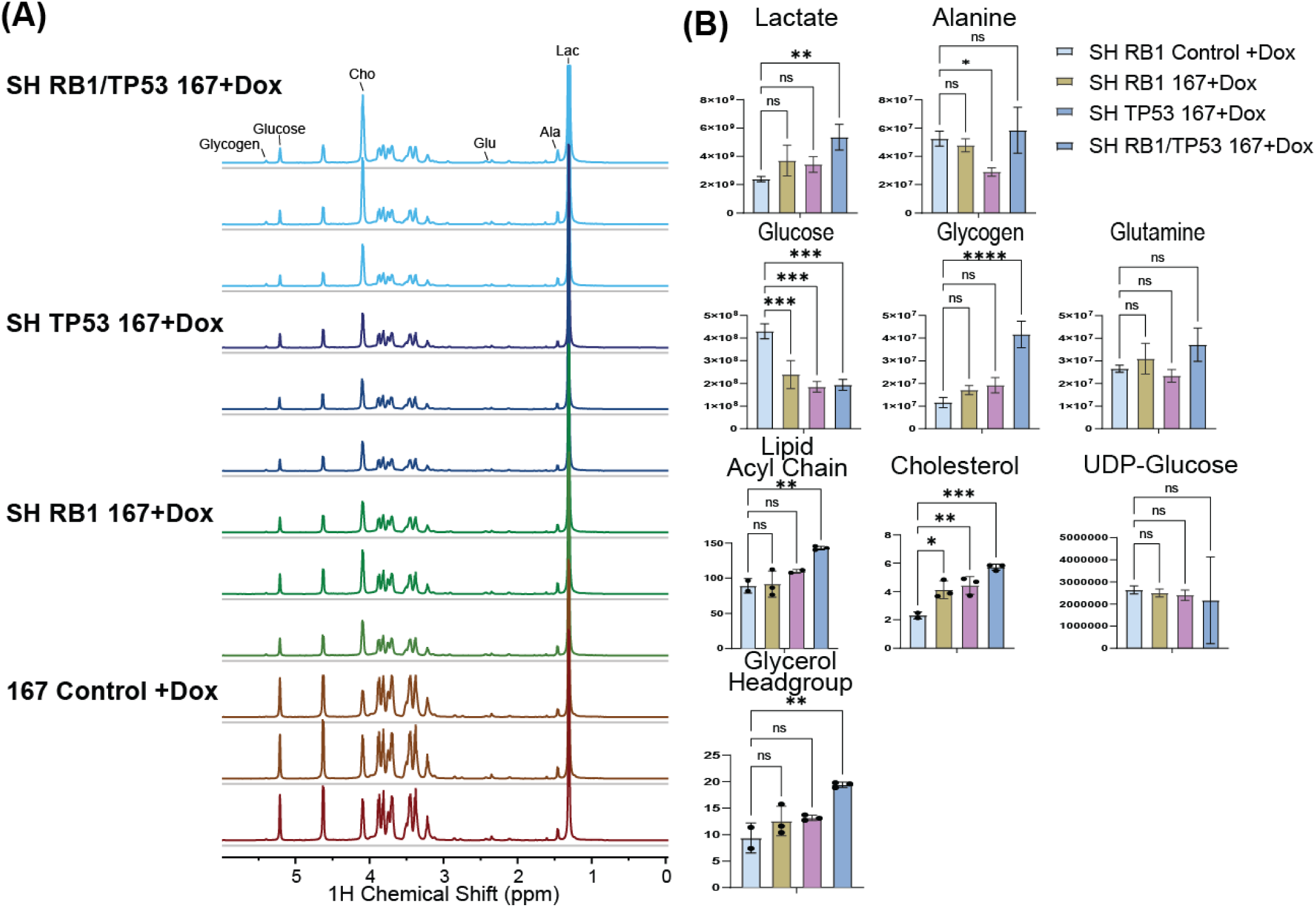
**(A)** ^1^H-^13^C HSQC NMR spectra of the polar fraction of 2 million LuCaP 167 organoid cells. Peaks used for assignment of specific metabolites are labeled by arrows **(B)** Quantification of metabolites from the polar and non-polar fractions. Multiplicity corrected p-values are calculated form two-way ANOVA test with Tukey’s correction for multiple comparisons using Prism 9.0. p= * (0.05), ** (0.01), *** (0.001). Error bars indicate SD.

**Supplementary Figure 6.**
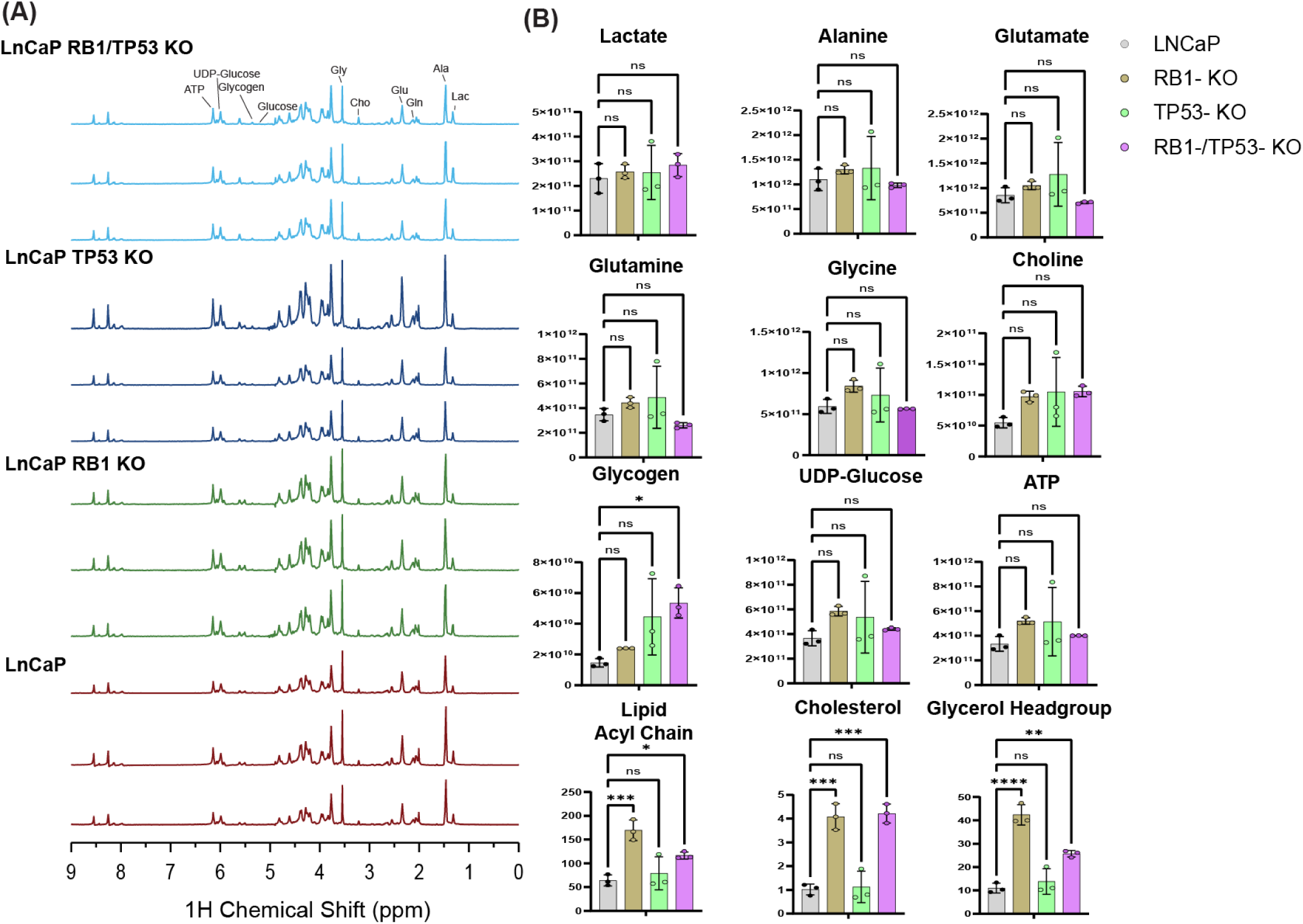
**(A)** ^1^H-^13^C HSQC NMR spectra of the polar fraction of LNCaP monoclonal Crispr KOs cells normalized to protein concentration by BCA. Peaks used for assignment of specific metabolites are labeled by arrows Quantification of metabolites from the polar and non-polar fractions. Multiplicity corrected p-values are calculated form two-way ANOVA test with Dunnet’s correction for multiple comparisons using Prism 9.0. p= * (0.05), ** (0.01), *** (0.001). Error bars indicate SD.

**Supplementary Figure 7.**
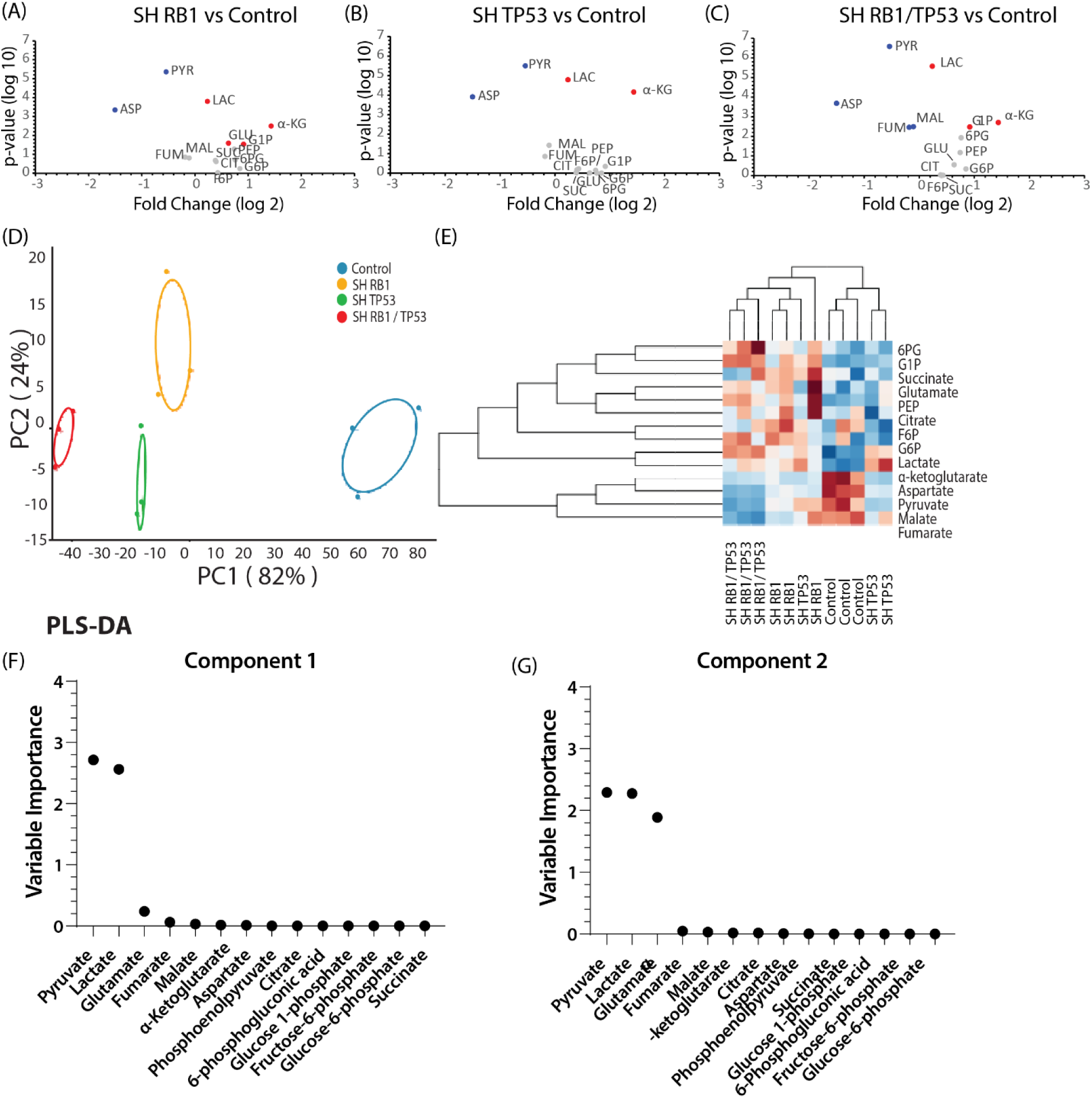
Metabolite concentrations from IC-MS of LuCaP 167 organoids with shRNA knockdown of *RB1* and/ or *TP53* relative to controls without shRNA. **(A to C)** Volcano plot of the log 2-fold change versus the associated p-value. Blue is significantly upregulated; red is significantly upregulated with **(D)** Principal component plot of the concentrations normalized to the total metabolite sum. The primary separation is the control samples from the RB1/TP53 depleted. **(E)** Heat map with hierarchal clustering **(F** and **G)** Variable importance for the first two components in a PLS-DA classification model for the four samples. The normalized concentrations of lactate, pyruvate, and fumarate are sufficient to separate the four samples (R^2^=0.572).

**Supplementary Figure 8.**
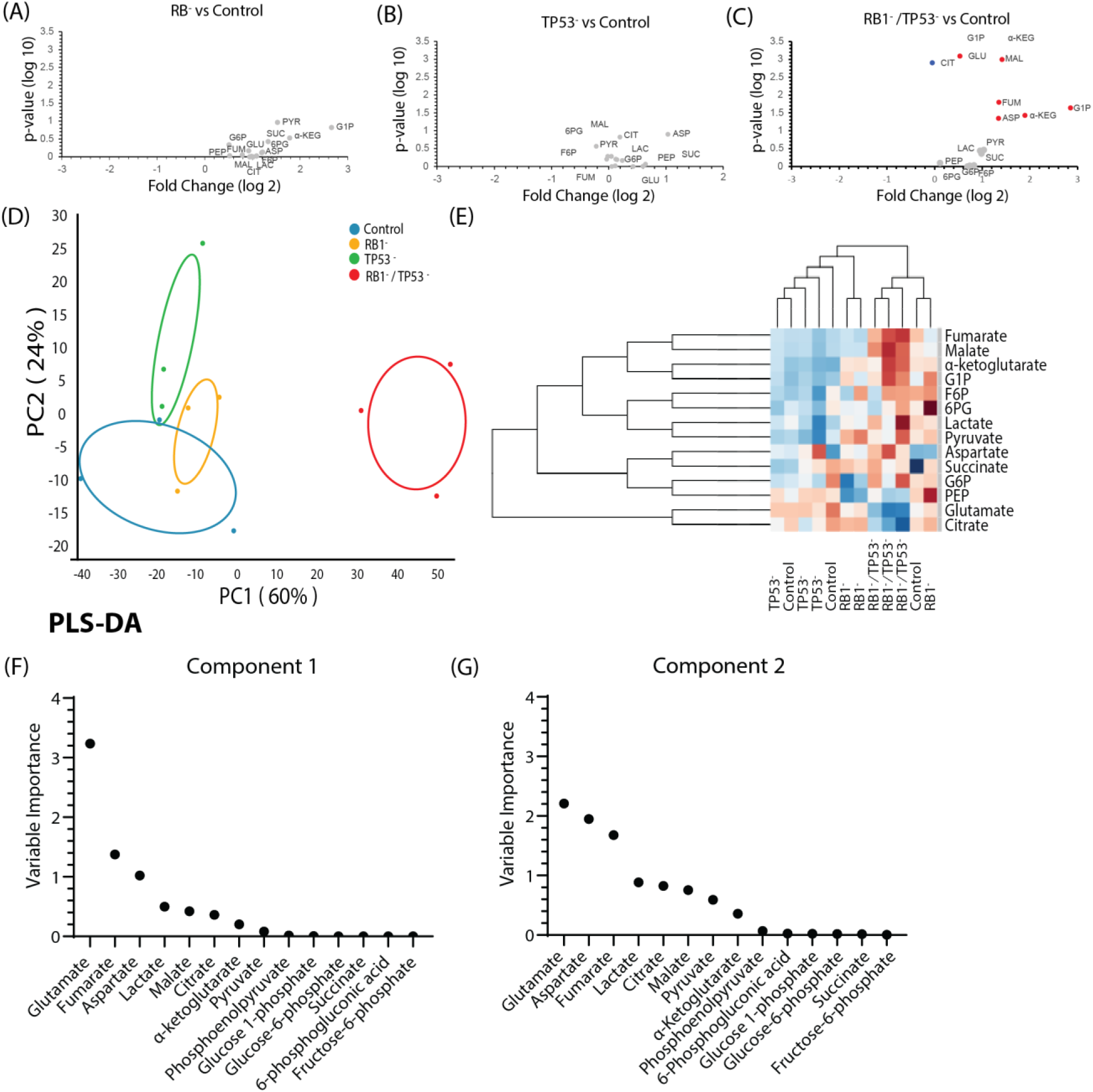
Metabolite concentrations from IC-MS of LnCaP monoclonal Crispr KO cells relative to control. **(A to C)** Volcano plot of the log 2-fold change versus the associated p-value. Blue is significantly upregulated; red is significantly upregulated with **(D)** Principal component plot of the normalized concentrations. The primary separation is the RB1/TP53-dual knockout from the others. **(E)** Heat map with hierarchal clustering **(F and G)** Variable importance for the first two components in a PLS-DA classification model for the four samples. The metabolites in the TCA cycle primarily separate the samples.

**Supplementary Figure 9.**
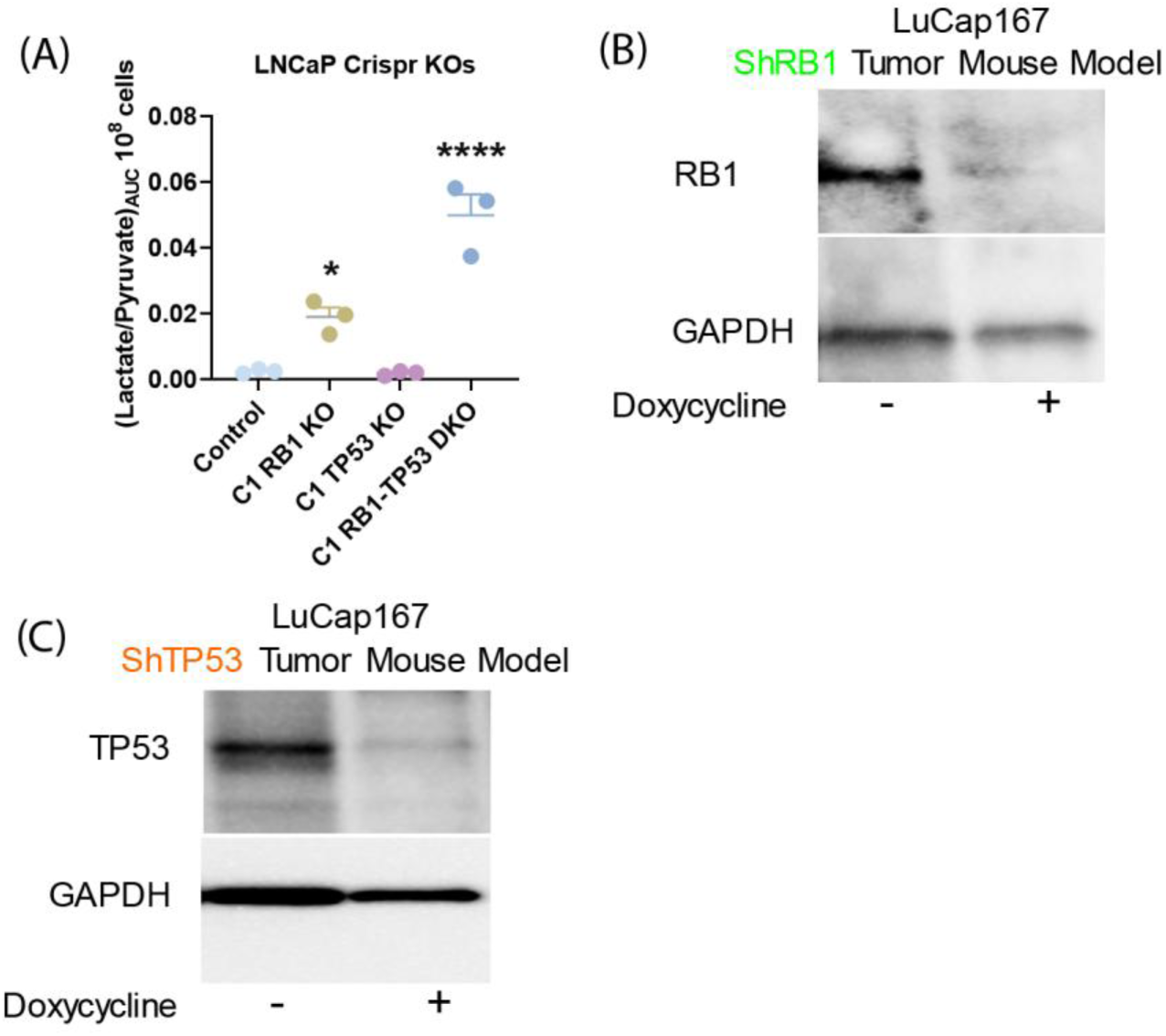
*RB1* alone or combined with *TP53* knockdown increases LDH flux detected *in vitro* by ^13^C-NMRS in LnCaP. **(A)** Dot plot of the lactate/pyruvate conversion ratio for genetically modified *RB1* and/or *TP53* LNCaP Crispr knockout models in vitro. **(B and C)** Western blot of RB1 (B) and TP53 (C) protein levels in LuCaP167 organoid-derived PDX tumors with or without *in vivo* doxycycline induction of ShRNA as described in methods.

## Supplementary Materials

The following supporting information can be downloaded at: www.mdpi.com/xxx/s1, Figure S1: Adenocarcinoma, castration-resistant prostate cancer models display higher FDG uptake; Figure S2: *RB1* and/ or *TP53* depletion maintains AR expression; Figure S3: *RB1* and/ or*TP53* knockdown does not affect: ^18^FDG^+^ uptake in LnCaP monoclonal Crispr knockouts; Figure S4: *RB1* and/ or*TP53* loss does not affect ^18^FDG^+^ uptake in a LuCaP PDX mode; Figure S5: ^1^H-^13^C HSQC NMR spectra of the polar fraction of 2 million LuCaP 167 organoid cells; Figure S6: ^1^H-^13^C HSQC NMR spectra of the polar fraction of LNCaP monoclonal Crispr KOs cells; Figure S7: Metabolite concentrations from IC-MS of LuCaP 167 organoids with shRNA knockdown of *RB1* and/ or *TP53*; Figure S8: Metabolite concentrations from IC-MS of LnCaP monoclonal Crispr KO cells relative to control. Figure S9: RB1 alone or combined with TP53 knockdown increases LDH flux detected in vitro by 13C-NMRS in LnCaP monoclonal Crispr Knockouts

## Author Contributions

Conceptualization, Kathleen Kelly; Formal analysis, Jeffrey Brender; Funding acquisition, Murali Cherukuri, Peter Choyke and Kathleen Kelly; Investigation, Fahim Ahmad, Margaret White, Kazutoshi Yamamoto, Daniel Crooks, Supreet Agarwal, Ye Yang, Brian Capaldo, Sonam Raj, Aian Alilin, Anita Ton, Stephen Adler, Jurgen Seidel, Colleen Olkowski and Jeffrey Brender; Methodology, Fahim Ahmad, Margaret White, Daniel Crooks, Ye Yang and Kathleen Kelly; Project administration, Murali Cherukuri, Peter Choyke, Kathleen Kelly and Jeffrey Brender; Resources, Kathleen Kelly; Supervision, Kazutoshi Yamamoto, Daniel Crooks, Murali Cherukuri and Peter Choyke; Writing – original draft, Fahim Ahmad and Jeffrey Brender; Writing – review & editing, Daniel Crooks, Ye Yang, Peter Choyke, Kathleen Kelly and Jeffrey Brender.

## Institutional Review Board Statement

The animal study protocol was approved by the Institutional Review Board of the Center for Cancer Research of the National Cancer Institute (protocols LGCP-003-2-C and RBB-159-3-P).

## Informed Consent Statement

Not applicable.

## Data Availability Statement

Dataset available on request from the authors

## Abbreviations

The following abbreviations are used in this manuscript:

AR: Androgen receptor
CR: Castration-resistant
CRPC: Castration resistant prostate cancer
NEPC: Neuroendocrine prostate cancer
ARPC: Androgen receptor positive adenocarcinoma
HP-MRI: Hyperpolarized magnetic resonance imaging
LDH: Lactate dehydrogenase
PDX: Patient derived xenograft
PC: Principal component
PLS-DA: Partial least-squares discriminant analysis
Rb1: Retinoblastoma 1 gene
TP53: Tumor protein 53 gene
18F-FDG: 18F-Fluorodeoxyglucose
SUV: Standardized Uptake Value
PSMA: Prostate Specific Membrane Antigen
VIP: Variable Importance in Projection

